# An *in situ* quantitative map of initial human colorectal HIV transmission

**DOI:** 10.1101/2022.04.30.490175

**Authors:** Heeva Baharlou, Nicolas Canete, Erica E Vine, Kevin Hu, Di Yuan, Kerrie J Sandgren, Kirstie M Bertram, Najla Nasr, Jake W Rhodes, Martijn P Gosselink, Angelina Di Re, Faizur Reza, Grahame Ctercteko, Nimalan Pathma-Nathan, Geoff Collins, James Toh, Ellis Patrick, Muzlifah A Haniffa, Jacob D. Estes, Scott N Byrne, Anthony L Cunningham, Andrew N Harman

## Abstract

The initial immune response to HIV is critical in determining transmission. However, due to technical limitations we still do not have a comparative map of early mucosal transmission events. We combined RNAscope, cyclic-immunofluorescence and novel image analysis tools to quantify HIV transmission dynamics in intact human colorectal tissue. We mapped HIV enrichment to mucosal dendritic cells (DC) and submucosal macrophages, but not CD4+ T-cells, the primary targets of downstream infection. DCs appeared to funnel virus to lymphoid aggregates which acted as early sanctuaries of high viral titres whilst facilitating HIV passage to the submucosa. Finally, HIV entry induced rapid recruitment and clustering of target cells, facilitating DC and macrophage mediated HIV transfer and enhanced infection of CD4+ T-cells. These data demonstrate a rapid response to HIV structured to maximise the likelihood of mucosal infection, and provide a framework for *in situ* studies of host pathogen interactions and immune mediated pathologies.

**Highlights:**

- *in situ* quantification of host cellular microenvironment response to pathogen invasion in human colorectal tissue.
- HIV first localises to mucosal DCs and submucosal macrophages, but not CD4+ T cells.
- Viral enrichment first occurs in lymphoid aggregates which is associated with passage into the submucosa.
- Early localisation of HIV to CD4+ T cells is associated with interactions with DCs and macrophages.

**Graphical Abstract:** 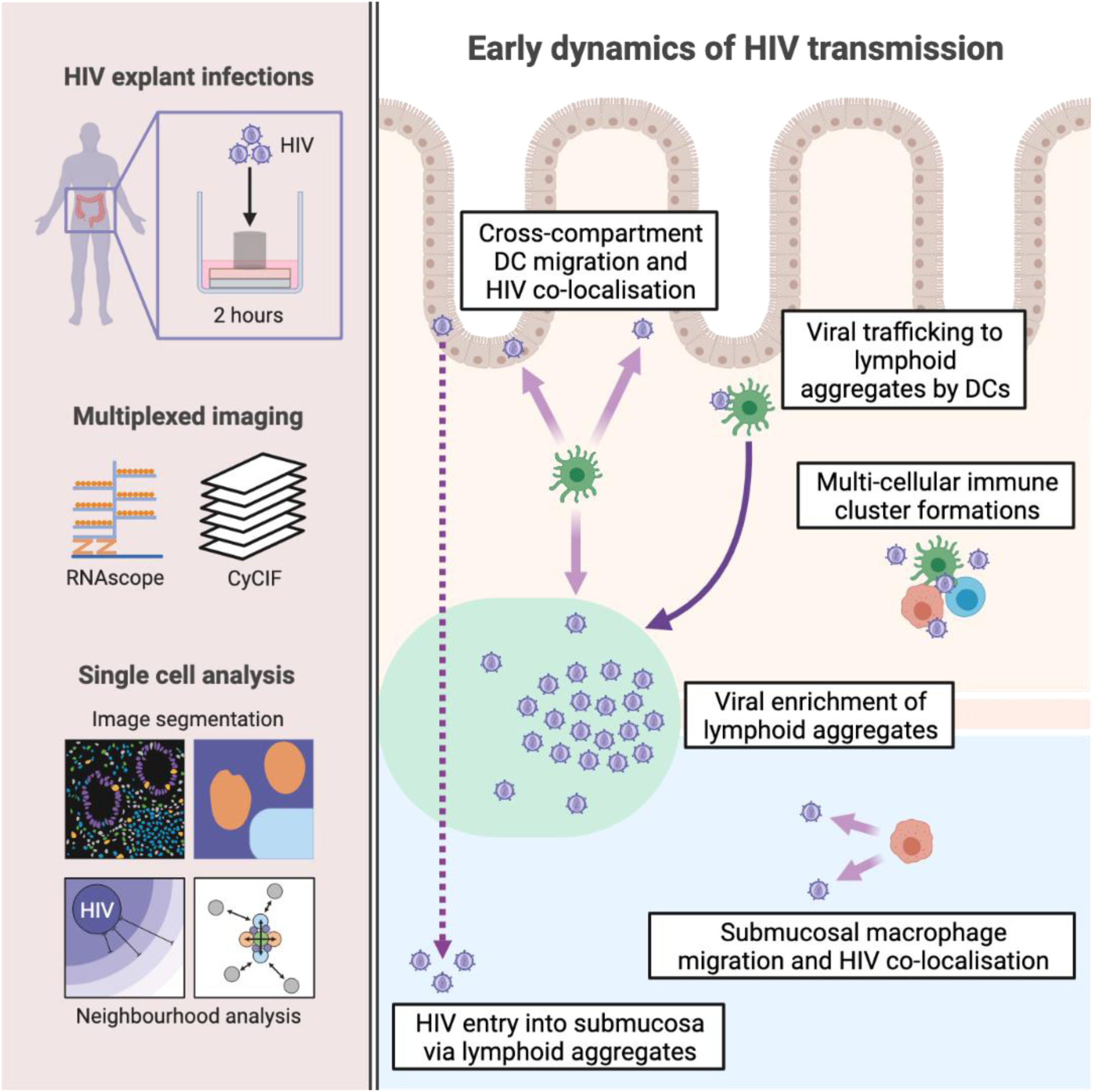

## Introduction

37 million people are infected with HIV and despite the introduction of pre-exposure prophylaxis there were still 1.5 million new infections in 2020. Blocking transmission of HIV therefore remains and high global priority which requires an effective vaccine and, in the meantime, better prophylactic interventions. To achieve these goals, we need a better understanding of the initial events that govern transmission of HIV particularly early viral interactions with antigen presenting cells such as dendritic cells (DC) and macrophages and their subsequent delivery of the virus to CD4+ T cells (Vine et al., 2022).

SIV challenge studies have shown that productive mucosal infection at the site of transmission precedes the detection of plasma viremia and that the viral reservoir is rapidly seeded at this site within days of exposure (Deleage et al., 2019; Whitney et al., 2014). Concordant data has also been described in human studies (Chun et al., 1998; Colby et al., 2018). Due to technological limitations early transmission studies have been limited to time points after initial viral integration/replication (Maric et al., 2021; Stieh et al., 2016; Vine et al., 2022) or the use of model systems, with human tissue studies mostly performed on isolated cells or by imaging of limited parameters (Bertram et al., 2019a; Ganor et al., 2010; Hladik et al., 2007; Rhodes et al., 2021; Trifonova et al., 2018; Vine et al., 2022).Thus, we still do not know the initial events that lead to mucosal HIV infection in the human tissues where transmission occurs.

The next stage in advancing our understanding of these events requires an *in situ* quantitative multi-parameter study to understand the relative involvement of multiple target cells within anatomically distinct tissue compartments. This ‘top down’ approach is critical for establishing physiological relevance and guiding the rational selection of specific biological mechanisms for in-depth characterisation studies. To our knowledge no study has examined all these processes at once in the context of pathogen invasion of human tissue. Such studies have been hampered to date by a plethora of issues including parameter limitations of conventional microscopy, a lack of suitable image processing and analysis algorithms and, in the context of HIV, difficulties with *in situ* pathogen detection at early time points (Deleage et al., 2016).

We have recently pioneered the use of RNAscope to visualise clinically relevant HIV virions interacting with anogenital target cells *in situ* within 2h of topical exposure (Bertram et al., 2019a; Rhodes et al., 2021). We have also designed post image acquisition algorithms to remove autofluorescence (Baharlou et al., 2020) and quantify cell interaction changes between disease states (Canete et al., 2022). In this study we have utilised these approaches as well as designed new tools to segment full cell bodies more accurately, allowing us to quantify the dynamics of HIV transmission across human colorectal tissue within 2h of exposure. We have defined the spatial distribution of the three key colorectal HIV target cells (DCs, macrophages and CD4+ T cells) across all colorectal tissue compartments (epithelium (EP), lamina propria (LP), lymphoid aggregates (LA) and submucosa (SM)) and shown how they respond to HIV. We show conclusively that HIV is first localised within mucosal (EP, LP, LA) DCs and submucosal macrophages, but not CD4+ T cells and that the virus is preferentially enriched in LAs. We also provide strong circumstantial evidence that LP DCs traffic virus to LAs and that these structures themselves may provide a conduit for rapid HIV entry into the deeper submucosal layer, where it preferentially associates with macrophages. Finally, we show that HIV mucosal entry induces its target cells to form multi-cellular clusters within which HIV+ DCs and macrophages preferentially cluster with CD4+ T cells, leading to viral transfer and enhanced infection of CD4+ T cells, supported by *ex vivo* data.

## Results

### Analysis pipeline for mapping HIV-target cell interactions *in situ*

This study explores the interactions of HIV with its three key target cells – DCs, macrophages and CD4+ T cells - in human colorectal tissue at the very earliest time points following HIV challenge using a combination of RNAscope, multiplexed fluorescence microscopy and novel analytical approaches (**Figure 1**). Lab adapted and transmitted/founder strains of HIV were applied to the apical surface of intact fresh human colorectal tissue for 2h. RNAscope was performed to detect HIV RNA (virions), followed by cyclic immunofluorescence (CyCIF) microscopy, which was used to identify nuclei (DAPI), EP (E-Cadherin), CD4+ T cells (CD3^+^CD4^+^), CD4- T cells (CD3^+^CD4^-^), DCs (CD11c^+^) and macrophages (FXIIIa^+^) (**Figure 1A-B**). While having little to no capacity for HIV uptake themselves, CD4- T cells (primarily CD8+ T cells) served as a useful control for non-specific HIV-cell interactions *in situ*. Post-acquisition, we removed autofluorescence signal, a known feature of colorectal tissue imaging (Wizenty et al., 2018) post-acquisition using our recently developed ‘Autofluorescence Identifier’ (AFid) algorithm (Baharlou et al., 2020) (**Figures 1A** and **S1A**). We next performed segmentation of HIV virions, cells, and tissue compartments (EP, LP, LA, SM) (**Figure 1C**). HIV RNA spots were segmented using a spot-segmentation algorithm (Battich et al., 2013). Cells were segmented using a multi-step approach whereby nuclei were first segmented, followed by manual cell type classification by marker expression, and finally cell body expansion. Importantly, cell bodies were expanded from individual nuclei in a manner that allows two cells to overlap in the same space, allowing for HIV virions to be accurately assigned to amorphous cells such as macrophages and DCs (**Figure S1B** and **S1C)**. This pipeline allowed us to make compartment specific measurements of target cell composition (**Figure 1D**), HIV status (HIV^+^ vs HIV^-^), virion load per cell (**Figure 1E**), target cell migration toward HIV (**Figure 1F**) and the effect of HIV on interactions between target cells (**Figure 1G**).

**Figure 1:**
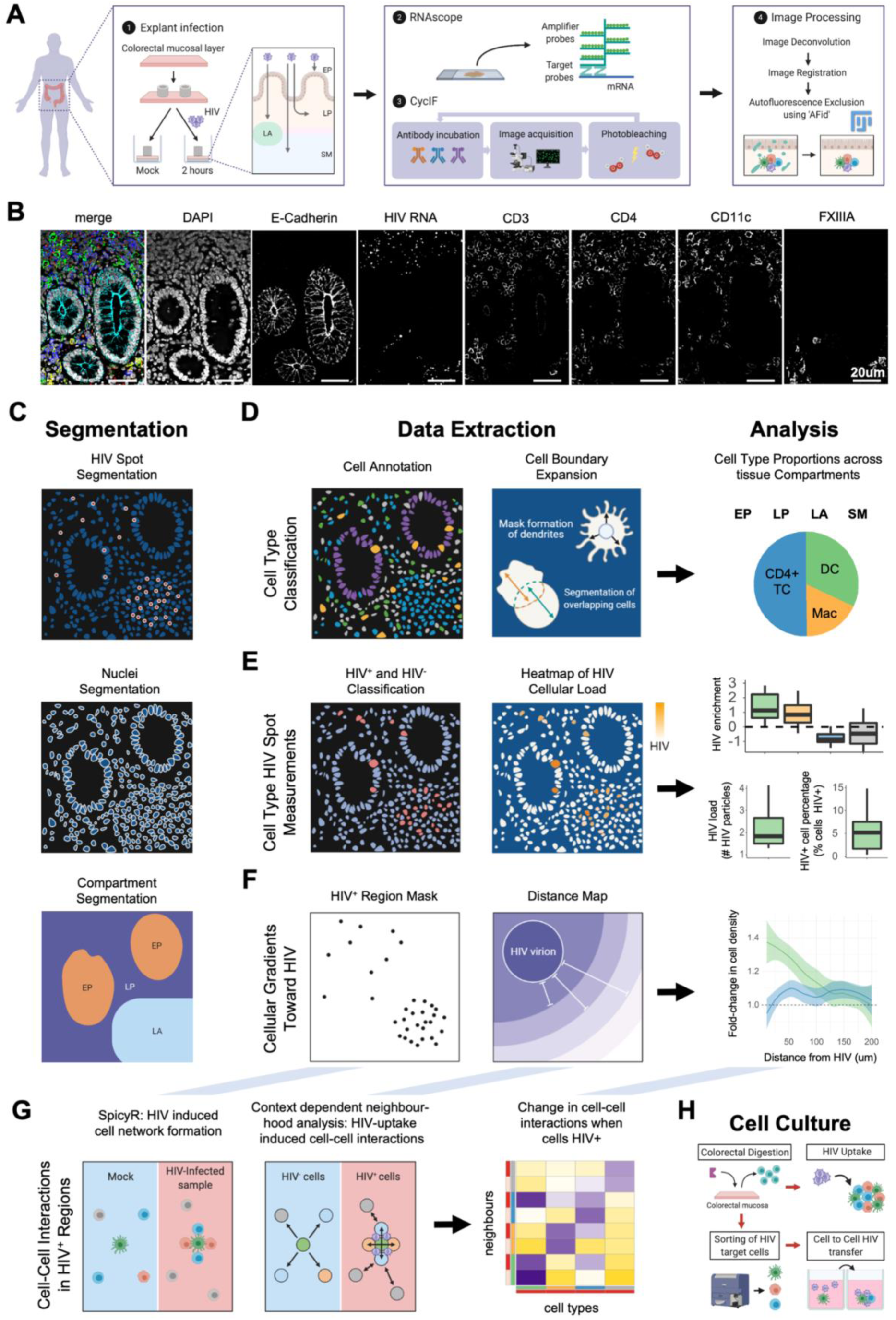
Analysis Pipeline for Mapping HIV-target Cell Interactions In Situ. **(A)** Explant infection and staining protocol. Within 15 minutes of surgery, human colorectal tissues were topically exposed to lab adapted HIV_Bal_ or clinical founder HIV_Z3678M_ for 2h, fixed, embedded and sectioned in transverse. 4um paraffin sections were then probed for HIV RNA using RNAscope, followed by CycIF staining to define key HIV target cells. Autofluorescence was then removed from the images post-acquisition using our Autofluorescence removal software ‘AFid’ (Baharlou et al., 2020). Images before and after Autofluorescence removal are shown in Figure S1A. **(B)** Representative images of markers acquired by RNAscope (HIV RNA) and CycIF (all remaining markers). **(C)** Segmentation was performed to estimate individual HIV virions, cell nuclei and the boundaries separating anatomical compartments of the colorectum (EP, LP, LA and SM) (SM compartment not shown in schematic). **(D)** Classification of HIV target cell types using a custom segmentation and classification approach outlined in detail in Figure S1C. Briefly, DCs, macrophages and T cells were classified using the percentage overlap of masks of markers CD11c, FXIIIa and CD3 with the segmented nuclei from part C. Cell bodies were then estimated using a custom approach that expands out from nuclei centers so as to estimate the actual cell boundary, regardless of cell shape and overlap with other cell types. **(E)** The segmentation of discrete HIV spots allowed for binary classification of cell types into HIV+ and HIV- categories (left) and forms an intuitive measure of HIV load in individual cell types (center). This allowed for comparisons of HIV association with HIV target cell types (right). **(F)** The creation of distance maps emanating from HIV particles (left and center) allowed for the measurement of how target cell densities fluctuate toward HIV and whether gradients form which may be indicative of cell migration. **(G)** Cell-Cell localization measurements were used to determine whether cell-cell interactions differed in HIV+ regions of tissue (left) and whether the uptake of HIV virions by target cells lead to unique cell-cell interactions (center). These measurements are performed between pairs of cell types and can be measured by an association score using a heatmap for visualization (right). **(H)** Tissue processing and HIV infection assay setup using ex vivo isolated HIV target cells. Immune cells were extracted from human colorectal tissue via collagenase incubation followed by ficoll density gradient enrichment and selection for CD45+ immune cells as per (Doyle et al., 2021). One of two assays was then performed. HIV uptake by target cells was assessed by treating CD45-selected colorectal cells with HIV_Bal_ for 2h followed by cell type identification and p24 assessment. The effect of myeloid cells on HIV infection of CD4+ T cells was assessed by treating sorted cells with HIV_Z3678M_ for 2h followed by incubation for 72h to measure newly synthesized virions. This was performed on either for CD4+ T cells alone, or on CD4+ T cells co-cultured with either DCs or macrophages.

To complement and validate these *in situ* data we used an orthogonal approach of flow cytometry analysis of HIV target cells following their dissociation from tissue and infection with HIV (**Figure S1D-F**). Using our HIV p24 uptake assay, we compared HIV-uptake *in situ* to that of *ex vivo* dissociated cells. Additionally, HIV-induced cell:cell interactions observed *in situ* (**Figure 1G**) were further investigated by sorting and infecting co-cultures of these cells to determine whether their interaction lead to enhanced viral transfer and infection (**Figure 1H).** Together these approaches enabled us to create an *in situ* quantitative map of how HIV is distributed across colorectal target cells and tissue compartments, as well as the HIV-induced cell-cell interactions that occur in the mucosa within 2h post-exposure and prior to systemic viral spread.

### HIV-target cell composition and distribution within human colorectal tissue

Using our analysis pipeline, we defined the relative proportion and density of HIV target cells within EP, LP, LAs and SM in fresh uninfected human colorectal tissue (**Figure 2A** and **2B**). Although only 1% of EP cells were HIV target cells, DCs were the most abundant (58%), followed by CD4+ T cells (28%) then macrophages (14%). Of the LP cells, 11% were HIV target cells with CD4+ T cells being most abundant (50%) followed by DCs (32%) then macrophages (18%). In LAs 35% of cells were HIV target cells consisting of CD4+ T cells (57%) and DCs (43%) with negligible presence of macrophages. 5% of SM cells were HIV target cells, with macrophages being most abundant (56%) followed by CD4+ T cells (25%) and DCs (19%). Cell density measurements closely followed these trends for each tissue compartment. Thus, LAs and LP harboured the highest density of HIV target cells with DCs and CD4+ T cells dominating the mucosal layers and macrophages dominating the submucosal layer.

**Figure 2:**
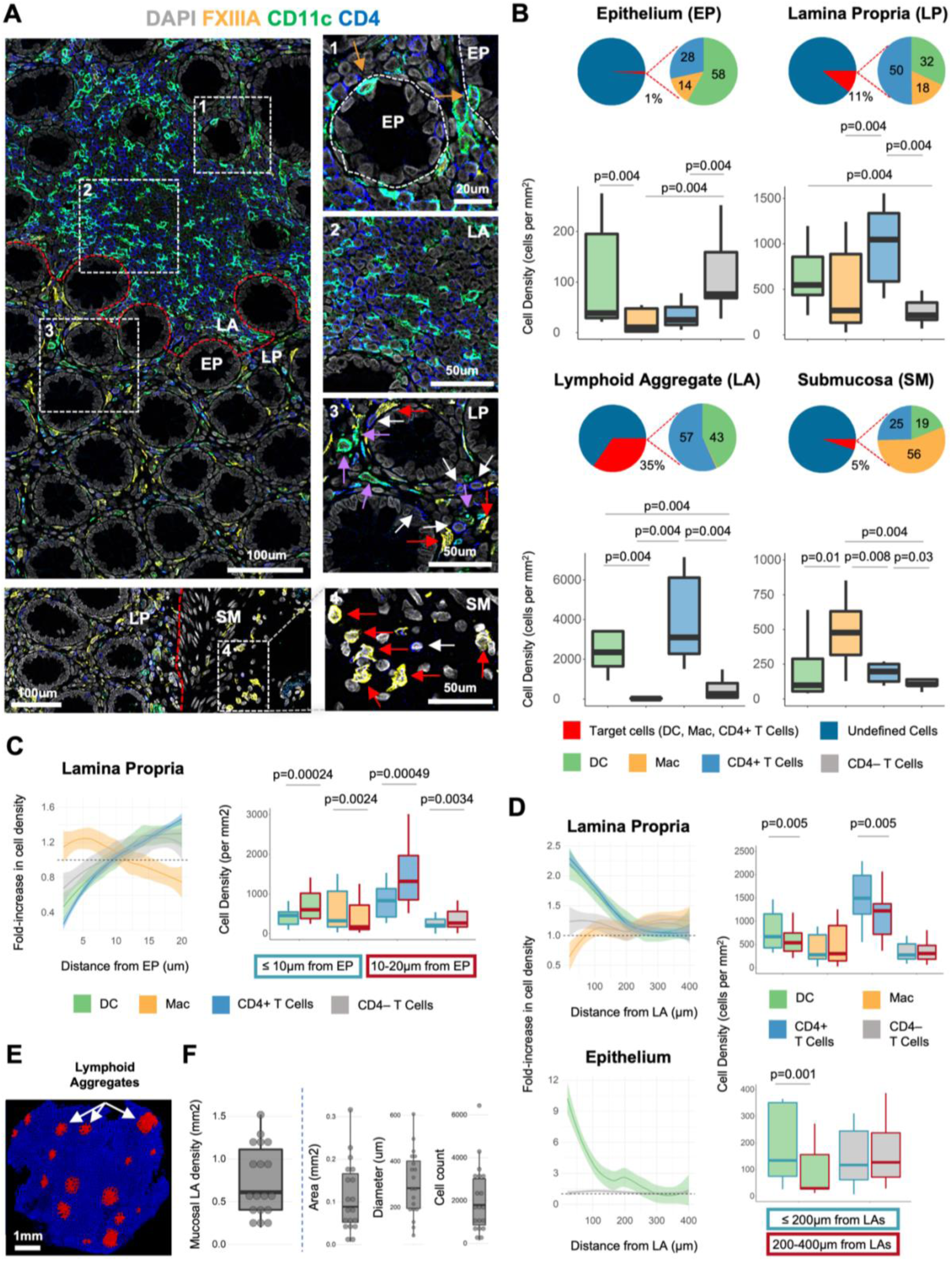
Distribution of HIV target cells across human colorectal tissue compartments. HIV target cell composition and spatial distributions within and across anatomical compartments of the colorectum were measured in uninfected healthy tissue. **(A)** Representative images of HIV target cells and their distribution in the human colorectum. A large field of view of colorectal tissue containing an LA is shown. (1) DCs in the EP (brown arrows) near the LA. (2) LA DCs and CD4+ T cells. (3) LP DCs (purple arrows), macrophages (red arrows) and CD4+ T cells (white arrows). (4) Submucosal macrophages (red arrows). The border between different tissue compartments is shown by the broken red line. The border of lymphoid aggregates was determined manually by the increased density of DCs and CD4+ T cells in these structures. **(B)** HIV target cell proportion and density across tissue compartments of the colorectum. Pie charts show the relative proportion of HIV target cells (DC, macrophage, CD4+ T cell) among all nucleated cells, as well as the composition of the HIV target cell population within each compartment. The density (per mm^2^) for target cells and CD4- T cells is shown below the pie charts for each tissue compartment. **(C)** Change in LP target cell density with distance from the EP. The left panel shows the fold-change in target cell density within 2µm non-cumulative intervals (0-2µm, 2-4µm… etc) from the EP compared to the target cell’s density in LP across the whole image. Each data point represents an individual donor (n=12). Results are shown as a LOESS curve of best fit to highlight LP cell density trends in relation to the EP. The right panel shows the density of HIV target cells and CD4- T cells in LP that are within 10µm (blue border) or 10-20µm (red border) from EP. **(D)** Change in EP or LP target cell density with distance from LAs. The left panel shows the fold-change in target cell density within 20µm non-cumulative intervals (0-20µm, 20-40µm… etc) from LAs vs the target cell’s compartment-specific density in the whole image. Each data point represents an individual donor where the results across LAs for a given donor were averaged (n=12). Results are shown as a LOESS curve of best fit to highlight cell density trends toward LAs. The right panel shows the density of HIV target cells and CD4- T cells in EP or LP that are within 200µm (blue border) or 200-400µm (red border) from LAs. **(E)** Representative image of the distribution of LAs in the human colorectal mucosa. The displayed image is the nuclei segmentation mask where LA-resident cells have been coloured red and all other cells blue. **(F)** Density (mm^2^), average area (mm^2^), diameter (µm) and cell count of LAs in the mucosa (EP + LP + LA) in 18 individual donors. Density measurements were performed as ‘cells per mm^2^ of DAPI’ for indicated regions. Statistical comparisons were performed using a Wilcoxon signed-rank test

We next examined the spatial distribution of HIV target cells within each compartment. This was achieved by using the border between tissue compartments as an anchor and measuring changes in cell density from this reference point. LP macrophages were enriched <10µm from the EP, whilst DCs, CD4+ T cells and CD4- T cells preferentially localised >10µm away from the EP (Wilcoxon, p<0.005 for all, **Figure 2C**). In LP, DCs and CD4+ T cells were enriched <200µm from LAs (Wilcoxon, p=0.005). This was also true for EP DCs (Wilcoxon, p=0.001) but was not measurable for EP CD4+ T cells due to their low frequency in EP (**Figures 2D**).

We next focused our attention on characterising LAs as they contained the highest density of HIV target cells, particularly DCs and CD4+ T cells (**Figure 2A-B**). LAs were present at a median density of 0.6 structures per mm^2^ of tissue and varied in their area (median 0.09mm^2^; range 0.05–0.17mm^2^), diameter (median 283µm; range 195 – 401µm) and cell number (median 1,800 cells; range 860 – 3015) (**Figure 2E-F)**.

### HIV viral particles are enriched in colorectal dendritic cells and macrophages within 2 hours

We next assessed the interactions between HIV and its target cells using lab adapted (BaL) or transmitted/founder (Z3678M) HIV strains. To ensure RNAscope probes were specific to HIV RNA we stained uninfected tissue and confirmed no signal was detectable (**Figure S2A**). As our pipeline incorporates automated virion detection, we compared mock and HIV-treated explants to calculate the false-detection rate. 1 particle per 1000 cells was detected in mock versus 30 particles per 1000 cells in HIV-treated explants, indicating only 3% of HIV+ cells in our treated samples were false positives (**Figure S2B**). To determine whether HIV was enriched among its target cells we measured the percentage of total HIV particles in this population (HIV Percentage), as well as the percentage of all cells that were target cells (Target Cell Percentage). A Chi-square test was then performed to test for whether enrichment among target cells was significant. The formula ‘log2(HIV Percentage/Target Cell Percentage)’ measured the relative level of HIV enrichment between images. Although the percentage of total HIV particles localising to target cells varied across images, the majority showed significant HIV enrichment in target cells compared to the remaining undefined ‘other cells’ (X^2^, p < 0.05, **Figure 3A**), with up to 4-fold enrichment observed in some images (**Figure 3A, bottom**). Visual inspection confirmed the specificity of HIV association with target cells over ‘other cells’ for HIV_BaL_ (**Figure 3B**) and HIV_Z3678M_ (**Figure S2B)**. Using a similar approach, we tested for HIV enrichment in different target cell subsets and compared the degree of enrichment between populations (**Figure 3C**). As both HIV_BaL_ and HIV_Z3678M_ had similar trends in the degree of enrichment for each cell type (**Figure S2C),** we combined these data to increase statistical power. HIV was enriched among DCs and macrophages (Wilcoxon, p<0.01), with higher enrichment (though not significant) in DCs (**Figure 3C**). Although, HIV associated with CD4+ T cells, this occurred less than expected relative to their abundance (Wilcoxon, p<0.01). Additionally, CD4- T cells associated with HIV at the same frequency as their relative abundance which is expected as they do not express HIV-binding receptors. We next analysed HIV-load in individual cells in each population to determine the differential amount of virus associated with each type of target cell. Combining RNAscope, spot-segmentation (**Figure 1C**) and cell-segmentation with cell body-estimation (**Figure S1B-C**) enabled us to accurately measure single-particle differences between cells (**Figure 3D**). We pooled cells across all samples, stratified them by HIV particle count and measured the target cell composition within each group (**Figure 3E**). Among the 12,822 target cells analysed, the majority (87%) contained only 1-3 HIV particles. Despite being the most prevalent cell type, CD4+ T cells were under-represented in all groups, particularly those with higher HIV particles numbers. Indeed, the only HIV target cells we could detect with >8 HIV particles/cell were DCs and macrophages, but never CD4+ T cells. This partly explains their relative HIV enrichment compared to CD4+ T cells (**Figure 3C**).

**Figure 3:**
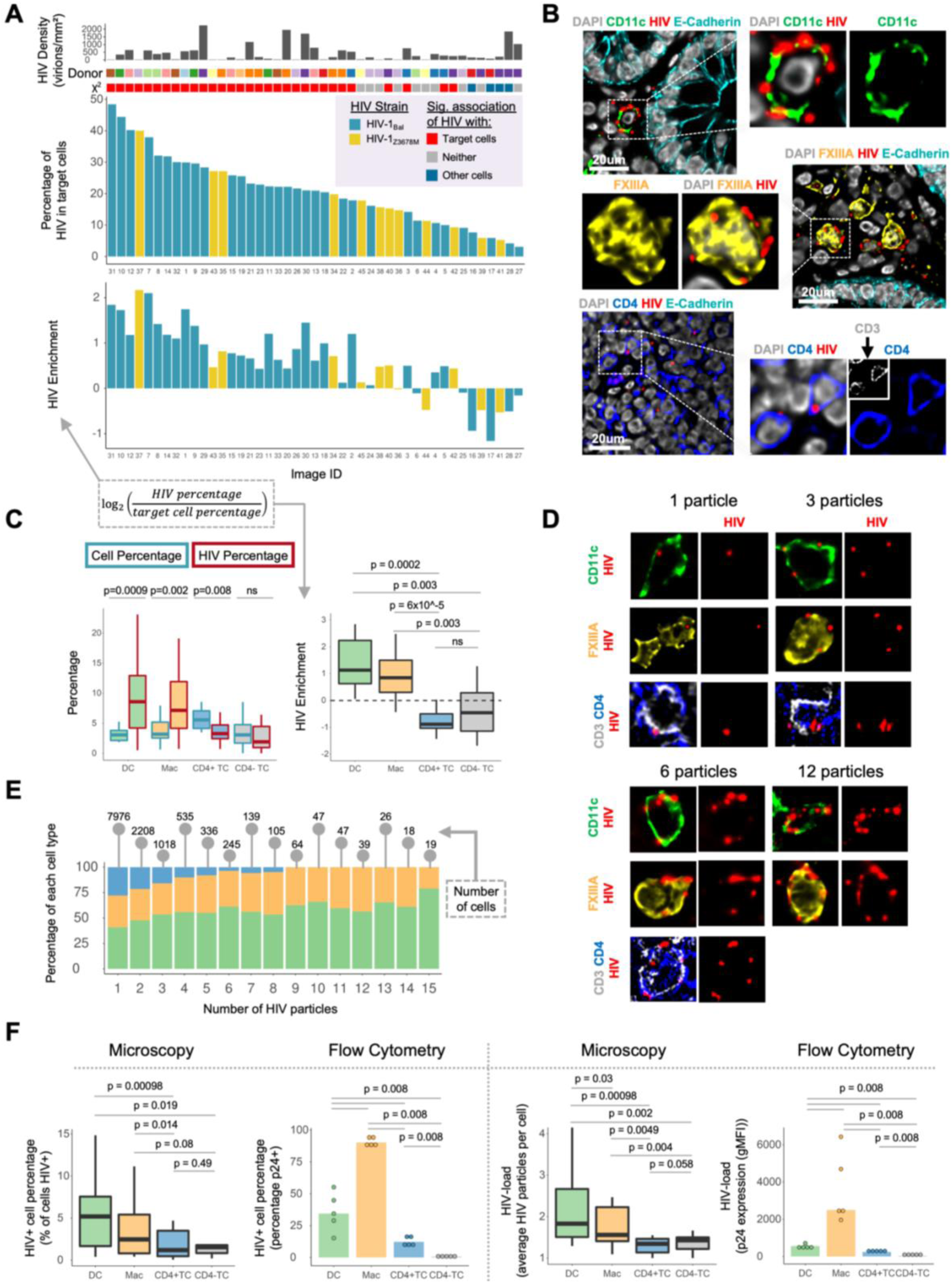
Assessment of interactions of HIV with colorectal target cells. **(A)** The middle graph shows the proportion of HIV particles (y axis) in individual images (x axis) associated with target cells (DCs, macrophages and CD4+ T cells) across 12 donors. Images are coloured by the HIV strain used (HIV_Bal_ or HIV_Z3678M_) for the explant infection. Top annotations for each image include the HIV particle density (mm^2^), donor number (color-coded) and outcome of a test for HIV-particle enrichment (Chi-square test (X^2^) of association; colours indicate significant enrichment of HIV in target cells (red), other cells (blue) or no significant enrichment in either population (grey); p < 0.05. Bottom annotation shows degree of HIV enrichment in the target cell population – calculated as log2(HIV percentage/target cell percentage), along with Pearson’s R and its p value for HIV enrichment vs HIV percentage. **(B)** Representative images of colorectal target cells interacting with HIV_Bal_ particles. **(C)** Comparison of HIV uptake (HIV Percentage) relative to opportunity (Cell Percentage) across target cells. For each cell type their percentage of all cells (blue border) or the percentage of all HIV particles in those cells (red border) is shown (left). A Wilcoxon signed-rank test was performed to test for HIV enrichment in each target cell population. The graph on the right shows the HIV particle percentage normalized to the cell percentage, and log2 transformed. A Wilcoxon signed-rank test was performed to compare this HIV enrichment score across cell types. Data points represent 15 explants from 12 donors, where the results from multiple images were averaged for each explant. **(D)** Representative images of target cells interacting with 1, 3, 6 or 12 HIV particles. **(E)** HIV+ target cells from all images were pooled and split into 15 groups depending on the number of HIV particles they contain (1-15 particles), and the composition of each cell type in each group was measured. The total number of cells in each group is annotated above each bar plot. **(F)** Comparison of uptake of HIV_Bal_ by target cells by Microscopy (left) and Flow Cytometry (right) in distinct donors. Left: The percentage of HIV+ cells which was defined as cells containing at least one HIV particle (microscopy) or cells with intracellular p24 expression (flow cytometry). Right: The average number of HIV particles per cell type (microscopy), or the mean fluorescence intensity of p24 staining in each cell type (flow cytometry). Statistical comparisons for microscopy measurements were performed using a Wilcoxon rank-sign test whilst flow cytometry measurements were compared using a Wilcoxon rank sum test. Microscopy: n = 11, Flow Cytometry: n =5. p < 0.01 = *, p < 0.001 = **, p < 0.0001 = ***; All density measurements were performed per mm^2^ of DAPI for indicated regions.

Finally, we sought to verify whether data derived *in situ* by microscopy would mirror that of *ex vivo* cells extracted from tissue using flow cytometry (**Figure 3F)**. This is important as the study of HIV transmission in human tissue has been largely confined to studies on *ex vivo* isolated cells (Ahmed et al., 2015; Bertram et al., 2019b; Botting et al., 2017; Rhodes et al., 2019). Both techniques confirmed that the ‘percentage of cells containing HIV’ and the ‘mean number of HIV particles per cell’ were significantly greater for DCs and macrophages compared to CD4+ T cells (Wilcoxon, p<0.01). However, flow cytometry measurements on *ex vivo* cells showed the CD4+ T cell ‘percentage’ and ‘average’ HIV measurements were significantly higher than CD4- T cells. In contrast no differences were observed by microscopy. This indicates that at early timepoints *in situ*, HIV interactions with CD4+ T cells are not yet measurably different to that of a random interaction between HIV and a cell type with no HIV- binding capacity. Furthermore, flow cytometry erroneously over-estimated macrophages as the dominant initial HIV-binding cell type, whereas *in situ* measurements showed DCs have a significantly higher mean HIV count per cell and a trend of higher HIV+ cell frequency (**Figure 3F**). These results highlight the importance of physiologically relevant quantitative *in situ* studies.

Taken together, these results demonstrate that HIV can enter human colorectal explants as early as 2h after inoculation resulting in preferential association with DCs and macrophages, compared to CD4+ T cells. Moreover, DCs and macrophages are capable of sampling larger quantities of virus at these early timepoints than other cells.

### HIV localisation patterns across colorectal tissue compartments and their associated target cells

We next turned our attention to tissue compartment specific differences in the distribution of HIV particles and HIV-containing target cells. LAs contained the highest density of HIV particles reaching over 10,000 virions/mm^2^ in several donors (Wilcoxon, p<3×10^-4^, **Figures 4A and S3A).** HIV density was lowest in the EP and SM with the density in LP higher than EP (Wilcoxon, p<0.03). To determine whether upon EP penetration, HIV preferentially localises to LP or LAs, we performed HIV enrichment analysis on compartments rather than cell types. In particular, we selected only LP and LA regions of images and compared the percentage of HIV in LAs to the expected percentage, defined as the percentage of the LP + LA area comprised of LAs (**Figure 4B**). Accordingly, values above or below expected (dotted line) indicate preferential localisation to LAs or LP respectively. This analysis revealed that HIV preferentially localised to LAs over LP (Wilcoxon, p=0.0007, **Figures 4C**), which was clearly observable upon visual inspection of images (**Figure 4D**).

**Figure 4:**
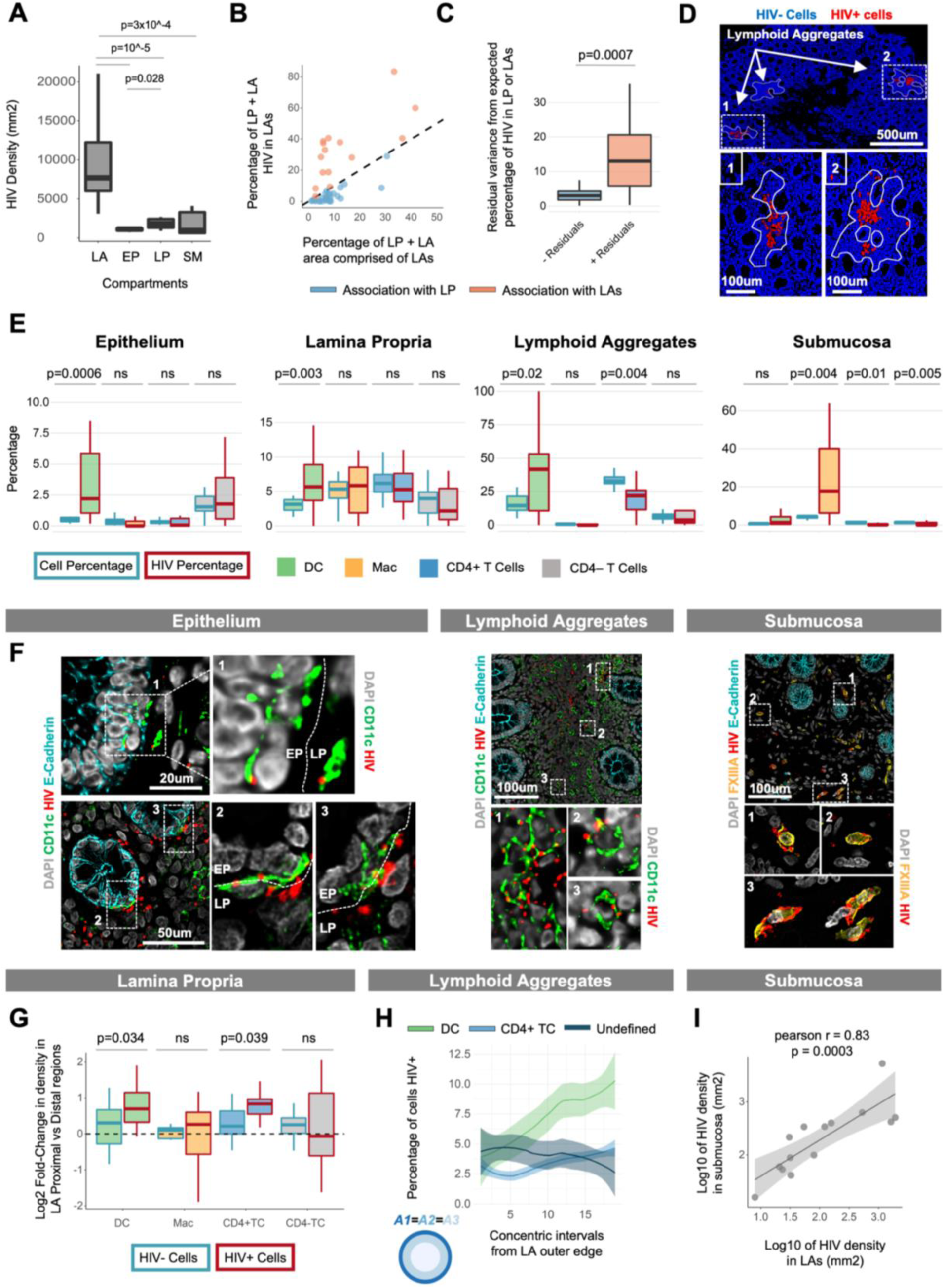
Differential HIV uptake across colorectal tissue compartments. **(A)** Comparison of HIV density (per mm^2^) across tissue compartments. Data for each compartment are the top 10 images with the highest HIV density for that compartment. A Wilcox rank-sum test was performed comparing HIV density across compartments. **(B)** Percentage of HIV virions (in the LP and LA) associated with LAs in each image vs the percentage of the image area (LP and LA only) comprised of LAs. The remaining percentages indicate values for the LP (e.g., rightmost data point is 60% LP which contains 40% of HIV). The dotted line is y=x and depicts the expected trend of data points in the case where there is no specific association or avoidance with either LP or LAs. Note that datapoints represent individual images across all donors. **(C)** Positive and negative residuals from (B) (perpendicular distance of points to the line y=x). A Wilcoxon rank-sum test was performed to compare the magnitude of difference between negative and positive residuals, which corresponds to preferential HIV localization to the LP and LAs respectively. **(D)** Representative images showing preferential localization of HIV to LAs compared to the surrounding LP. The displayed image is the nuclei segmentation mask showing HIV+ cells (red) and HIV- cells (blue). The border of lymphoid aggregates was determined manually by the increased density of DCs and CD4+ T cells in these structures. **(E)** Comparison of HIV uptake (HIV Percentage) relative to opportunity (Cell Percentage) for each cell type in different compartments of the colorectum. For each cell type their percentage among all cells (blue border) or the percentage of all HIV particles in the image in those cells (red border) is shown. A Wilcoxon signed-rank test was performed to test for HIV enrichment in each target cell population. **(F)** Representative images of DCs interacting with HIV in EP and LAs, as well as macrophage-HIV interactions in the SM. The broken line indicates the border between EP and LP. **(G)** Log2 fold-change in either HIV- or HIV+ cell density in LA-proximal (<=400µm) vs -distal (400-800µm) regions of the LP. A distance of 400µm was chosen (instead of 200µm as in Figure 2D) in order to capture a larger pool of HIV+ cells, thus improving the reliability of comparisons. Data represent the ‘proximal’ and ‘distal’ LP from individual LAs. Data was assessed for normality and a paired t-test was performed comparing cell density between regions. Paired regions for a given cell type were only considered if they each contained at least 5 HIV+ and 5 HIV- cells to allow for a fair comparison of density between regions. **(H)** Percentage of mucosal LA DCs, CD4+ T cells or undefined cells (not a DC, macrophage or T cell) that are HIV+ in non-cumulative intervals from the outer edge of HIV+ LAs (x = 1) toward their center (x = 20). Results are shown as a LOESS curve of best fit to highlight cell density trends from outside LAs toward their center. Statistical comparisons on discrete concentric intervals are shown in Figure S4G. **(I)** HIV virion density (per mm^2^ of DAPI) in SM vs LAs. Smoothed linear regression is shown with Pearson’s r and its associated p value. All density measurements were performed per mm^2^ of DAPI for indicated regions.

Having observed HIV enrichment in DCs and macrophages (**Figure 3C**), we next determined the specific compartments in which this viral enrichment occurred. In EP, LP and LAs HIV preferentially localised to DCs, whereas in SM the virus preferentially localised to macrophages (**Figures 4E).** Representative images are shown in **Figure 3B top panel** for LP and **Figure 4F** for all other compartments. In LAs 25% of donors showed >50% of HIV localised to DCs, which is far more than other compartments (**Figure 4E).** Correspondingly, LA CD4+ T cells harboured significantly less HIV than expected based on their frequency. Therefore, LA DCs may be more primed for antigen-uptake, which would partly explain the high levels of HIV observed in these cells in this compartment.

Comparing the degree of HIV enrichment in cells between compartments, we observed that EP DCs had a 2-fold HIV enrichment compared to LP and LA DCs. In contrast, SM macrophages had a median 4-fold enrichment compared to their LP counterparts (Wilcoxon, p<0.05, **Figure S3B**). These compartment specific differences in enrichment are explained by differences in cellular HIV-load and the percentage of the population interacting with HIV. In particular, EP DCs had a higher HIV-load per cell and a higher frequency of interactions with HIV, whereas SM macrophages had an increased frequency of HIV interactions, but a similar viral load to their LP counterparts (Wilcoxon, p<0.05, **Figure S3C-D**).

Beyond mapping HIV-target cell interactions to compartments we also explored how HIV+ cells were spatially distributed within compartments. Beginning with LP, we observed that like uninfected tissue (**Figure 2D**), HIV+ DCs and CD4+ T cells were enriched near LAs (**Figure S3E**). However, this enrichment was greater for HIV+ DCs and CD4+ T cells than their HIV- counterparts (Paired two-sample t-test, p<0.05, **Figure 4G**), despite no difference in the frequency of viral particles between LA- proximal or -distal regions of the LP (**Figure S3F)**. Interestingly, despite our previous observation of enrichment of both LP macrophages near EP and EP DCs near LAs (**Figure 2C-D**), we did not observe any enrichment of HIV+ populations of these cells near these structures (data not shown).

In LAs themselves we measured HIV particle distribution using the formula for even-area concentric rings to divide these structures into roughly equal area intervals (**Figure S3G**). Using this approach, we observed a significant increase in HIV density towards the centre of LAs (**Figure S3H).** Stratifying LAs by size **(Figure S3I**), HIV particles could even be detected toward the centre of larger LAs (>500µm in diameter), suggesting there may be a mechanism to focus virus centrally within these structures (**Figure S3J**). Measuring HIV+ cells rather than individual particles, we observed that DCs were the only HIV-containing cell type to increase significantly in frequency toward the centre of LAs (**Figures 4H** and **S3K**). This suggests that DC-mediated transport may in part contribute to the central focusing of HIV within LAs.

Finally, as we were surprised that HIV interacted with macrophages in the deep SM layer as early as 2h, we turned our attention to this compartment. We first confirmed that HIV entry into the SM correlated with entry into the overlying mucosal layer (Pearson R: 0.62, p=10^-5^, **Figure S3L**). Next, we fitted a linear model of HIV density in the SM as a function of HIV density in either the LP, LAs or both compartments (**Figure S3M**). Although both compartments significantly predicted HIV entry into the SM when measured in isolation, the combined model revealed LAs as the only significant predictor of SM HIV entry. This is likely explained by the high correlation between LP and LA HIV density (**Figure S3N, last row**) leading to confounding of the LP variable when modelling the determinants of SM HIV localisation. A scatterplot of SM vs LA HIV density, illustrates a clear correlation between these measurements (Pearson R: 0.83, p=0.0003, **Figure 4I**). Furthermore, we observed HIV particles from the LA apical surface within the mucosa, right through to the basal surface in the SM (**Figure S3N**). This suggests that viral trafficking through the length of LAs is possible.

Finally, we present multiple lines of evidence to confirm that SM entry was not due to leakage of virus from the cloning cylinder during the culture period. First, we observed a high correlation between mucosal and SM HIV densities (**Figure S3L**), suggesting ordered entry of HIV into the SM from the mucosal layer. Second, HIV+ macrophages were in both superficial (near crypt bases) and deeper regions of the SM (**Figure S3O**), whereas leakage would likely result in virus predominately in deeper regions, toward the bottom of the tissue. Finally, we collected the explant culture media at the end of the culture period and confirmed that no HIV was present using a sensitive HIV detection assay (**Figure S3P**).

Together these results reveal substantial differences in HIV distribution across colorectal tissue compartments, with LAs as key initial entry sites appearing to facilitate HIV access to the SM. Furthermore, HIV enrichment in DCs and macrophages occurred only in the mucosa and SM respectively. This highlights that it is not only the cell type, but also its compartmental residence which determines the degree of interaction with HIV.

### Colorectal dendritic cells migrate within and across tissue compartments in response to HIV

We next explored potential mechanisms of differential HIV-target cell interactions by analysing the spatial organisation of target cells in relation to HIV. In particular, we measured changes in target cell density from HIV particles, where steadily increasing or decreasing density gradients were used to infer cell migration in response to HIV (**Figure 1F**). In EP, LP and LAs we observed that DCs formed an increasing density gradient toward HIV, whereas T cells and macrophages did not (**Figure 5A**). The effect was most pronounced in LP to the extent that DCs were depleted in regions >300μm from HIV. Similarly, SM macrophages formed an increasing density gradient toward HIV (**Figure 5A**).

**Figure 5:**
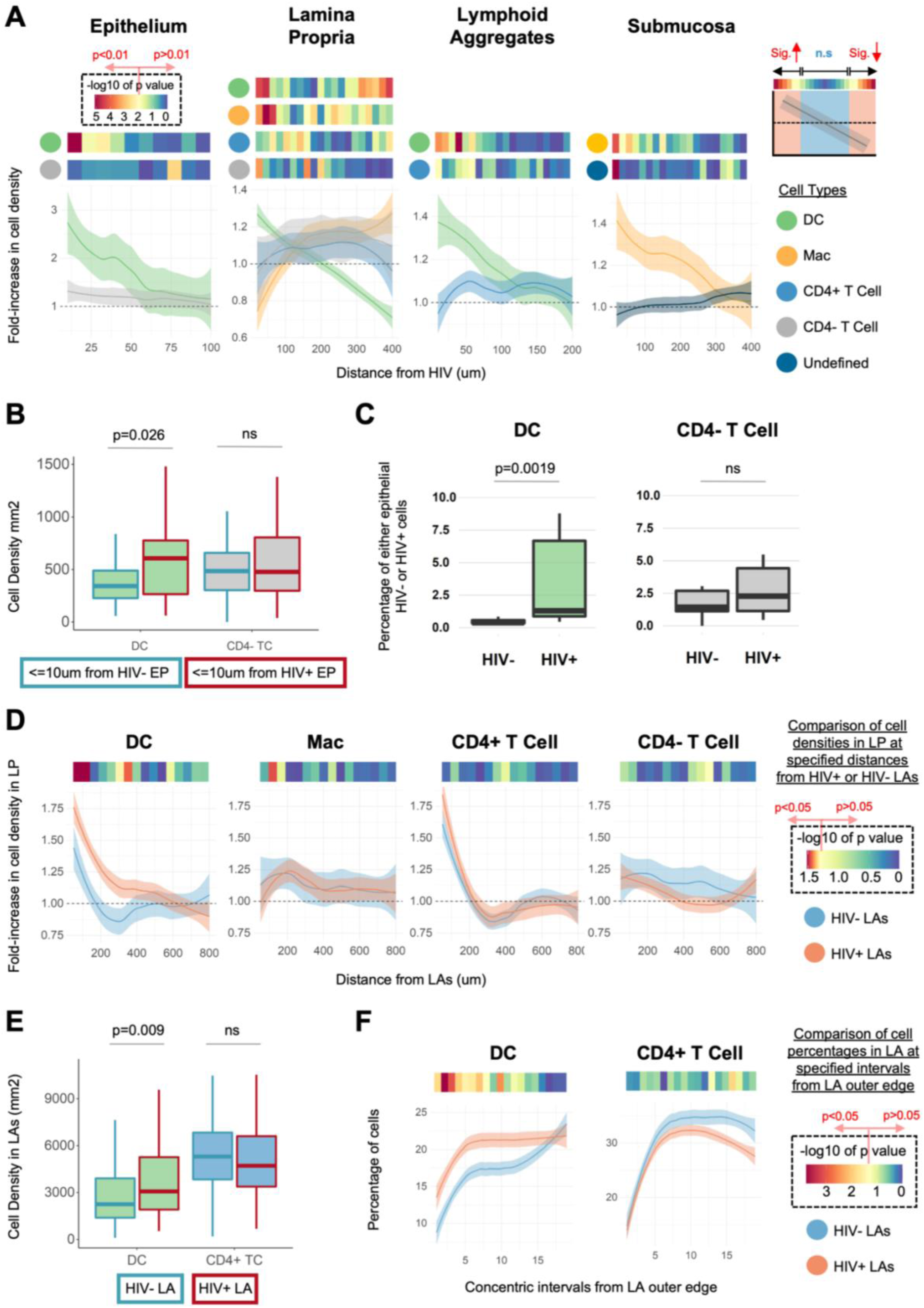
Cellular gradients in response to HIV. **(A)** Changes in target cell density with distance from HIV, measured in different tissue compartments. Measurements are the fold-change in target cell density within 10µm (EP) or 20µm (LP, LA, SM) non-cumulative intervals from HIV. Fold-change was measured in relation to the target cell’s average density in its tissue compartment. Each data point represents an individual image (n=45 in 12 donors). Results are shown as a LOESS curve of best fit to highlight cell density trends in relation to HIV. Statistical comparisons were performed between the target cell density in each interval and the average density of the target cell in the compartment (Wilcoxon signed-rank test). These are annotated above the graphs. P value heatmaps were centered at p=0.01 (cream colour) which was designated as the threshold for significant changes in cell density. **(B)** The density of LP DCs and CD4- T cells was measured within 10µm of HIV+ EP (containing at least 1 HIV particle) or HIV- EP (at least 50µm away from HIV+ EP). Only images with at least one cell in each category were analysed. Images were binned into 100×100µm quadrats and only quadrats with more HIV particles in EP than LP were kept for analysis. Data represents individual images (n=40 in 12 donors). A Wilcoxon signed-rank test was performed comparing cell densities near HIV+ EP vs HIV- EP for each image. **(C)** EP cells were classified as HIV+ or HIV- and the percentage of each population comprising DCs or CD4- T cells was measured. Data represent individual donors (n=12). A Wilcoxon signed-rank test was performed comparing cell percentages between HIV+ and HIV- epithelia. **(D)** Changes in target cell density with distance from HIV+ (>=2 particles) or HIV- (<2 particles) LAs. Measurements are the fold-change in target cell density within 50µm non-cumulative intervals from LAs. Fold-change was measured in relation to the target cell’s average density in the image. Data represent individual images (n=38 in 12 donors). Results are shown as a LOESS curve of best fit to highlight cell density trends in relation to LAs. Statistical comparisons were performed, comparing target cell densities in each interval from HIV+ vs HIV- LAs in the same image (Wilcoxon signed-rank test). These are annotated above the graphs. P value heatmaps were centered at p=0.05 (cream colour) which was designated as the threshold for significant differences in cell density. **(E)** Density of DCs or CD4+ T cells in mucosal HIV+ vs HIV- LAs. Only donors with both HIV+ and HIV- LAs were used in this analysis. Statistical comparisons were performed using a Wilcoxon rank-sum test to compare the magnitude of cell density changes between HIV- LAs (n=84) and HIV+ LAs (n=82) in 11 donors. **(F)** Comparison of DC or CD4+ T cell frequency (percentage of all cells) in non-cumulative intervals of mucosal HIV+ vs HIV- LAs from their outer edge of HIV+ LAs (x = 1) toward their center (x = 20). Results are shown as a LOESS curve of best fit to highlight cell density trends from outside LAs toward their center. Statistical comparisons were performed, comparing target cell frequencies in each interval between HIV+ and HIV- LAs (Wilcoxon rank-sum test). These are annotated above the graphs. P value heatmaps were centered at p=0.05 (cream colour) which was designated as the threshold for significant differences in cell density.

As DCs appeared to migrate toward HIV in all three mucosal compartments, we investigated whether they also could cross tissue compartments in response to HIV. To this end, we measured whether LP DCs redistributed ‘toward’ and ‘into’ EP or LAs when these compartments contained HIV. Starting with EP, we classified EP cells into HIV+ or HIV- populations and measured the density of DCs in the LP ≤10µm from each EP population. Compared to HIV- EP, DCs were significantly more concentrated beneath (**Figure 5B**) and within HIV+ EP (**Figure 5C**). Using a similar approach for LAs, we found that compared to HIV- LAs, LP DCs were significantly more enriched near HIV+ LAs **(Figure 5D**) and DCs were present at a higher density within HIV+ LAs (**Figure 5E**). Additionally, DCs were more concentrated toward the outer edge of HIV+ LAs, further supporting the idea that DCs migrate into these structures in response to HIV (**Figure 5F**). We did not observe any difference in CD4+ T cell density near or within HIV+ vs HIV- LAs.

Taken together, these results suggest that HIV enrichment in mucosal DCs and SM macrophages may relate to their ability to migrate toward incoming HIV, with DCs going as far as to cross tissue compartments in response to incoming virus.

### HIV induces the formation of target cell clusters within which dendritic cells and macrophages traffic virus to CD4+ T cells

As cell:cell interactions are important for the spread of HIV (Bertram et al., 2019b; Bracq et al., 2018) we performed spatial analysis to determine if HIV influenced interactions between target cells *in situ.* We used two different but complementary approaches, one developed by us called ‘SpicyR’ (Canete et al., 2022), and a modified approach to ‘Neighbourhood Analysis’ (Schapiro et al., 2017). We used SpicyR to compare cell:cell interactions between HIV+ and HIV- regions of treated explants. This allowed us to determine whether the presence of HIV influences cell:cell interactions, regardless of whether the cells themselves are HIV+. Within HIV+ regions, Neighbourhood Analysis then determined whether HIV-uptake itself induces cell:cell interactions.

SpicyR showed that in HIV+ regions, target cells cluster amongst each other in both LP and LAs. Interestingly, LP non-target CD4- T cells clustered amongst themselves in HIV+ regions but not with HIV target cells (**Figure 6A**). Additionally, there was no difference between HIV- regions and the mock explants, confirming that clustering occurs specifically in response to HIV (**Figure 6A**). To verify that results are resistant to parameter variation, we ran SpicyR over a range of radius and distance cut offs (**Figure S4A**). Qualitative inspection of images confirmed the presence of clusters of DCs, macrophages and CD4+ T cells in HIV+ regions (**Figure 6B**). Interestingly, clusters had a tendency to form away from the EP in the central area between the crypts of Lieberkühn. We hypothesised this was due to target cell migration away from the EP interface in response to incoming HIV. To investigate this, we devised a method of temporal inference to model the progression of HIV entry into the mucosa. Here, images were divided into 100µm^2^ windows, each classified as ‘naïve’ (HIV density = 0), ‘early’ (HIV density EP > LP) or ‘late’ (HIV density EP < LP) in terms of HIV entry, with changes in target cell density measured close and far from the EP interface at each stage. Results showed that like uninfected tissue, macrophages were enriched near the EP in naïve regions, however early in response to HIV they migrate deeper and persist there in the late phase (**Figure S4B**). Interestingly we found that CD4+ T cells also migrate away from the EP, but only in the late phase, whereas CD4- T cells showed no change throughout the phases of entry. Curiously, DCs also showed no difference, though we can presume this is due to their dual role of migration toward HIV+ EP (**Figure 5B-C**) as well as forming part of the central clusters in the LP (**Figure 6A and B**). These results suggest that a secondary effect of colorectal HIV entry is the rapid formation of a target-cell enriched community slightly distal to the site of entry, thus creating an ideal environment for cell-to-cell viral transfer.

**Figure 6:**
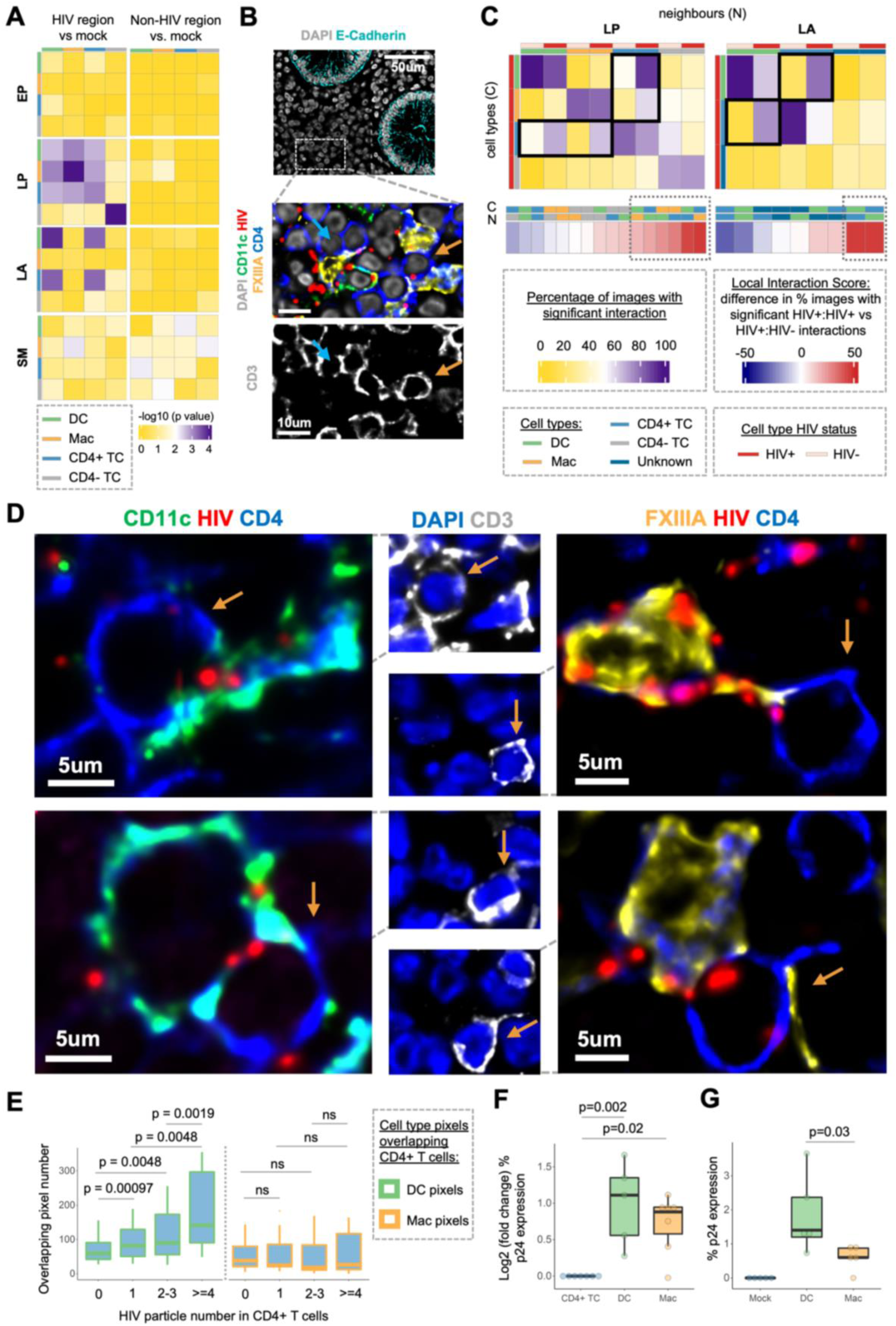
Dynamics of HIV-induced cell-cell interactions in situ. **(A)** SpicyR analysis to assess differential cell-cell localization between HIV and mock-treated samples in the EP, LP, LA and SM. The HIV-proximal region was defined as all cells within 30µm of HIV for the EP, LP and LA compartments, and 100µm for the SM due to the lower cell density in this compartment. As a control, HIV-distal regions (>30µm or 100µm) were compared to the mock sample. P values were centered at p=0.01 which was the assigned cut off for significant changes for this analysis. **(B)** Representative images of HIV target cell clustering in a HIV+ region located away from the EP interface. Brown arrows indicate clusters of target cells. **(C)** Neighbourhood analysis to assess HIV-uptake induced target cell interactions. HIV+ target cell neighbourhoods were measured by cell type and whether they were HIV+ or HIV-. Neighborhoods were defined as any cell within 6µm from the edge of a given cell. These neighbourhoods were then assessed for significance by comparison to the bootstrapped distribution of HIV+ cell labels in the image. The distribution was generated with 999 permutations and the cut off for significance defined as p<0.005. The percentage of images showing a significant interaction for each cell-cell pair is shown as a heatmap where values range from 0% to 100% of images. Also shown at the bottom is the ‘local interaction score’, defined as the percentage difference in images showing a significant interaction when a given neighbor is HIV+ vs HIV-. Cell types and their HIV status (HIV+ vs HIV-) are annotated and key interactions are encased by boxes. **(D)** Representative images of DCs and macrophages interacting with CD4+ T cells where HIVBal is present at the interface between cells. For clarity, CD3 staining is shown in a separate image with a brown arrow pointing to CD3+CD4+ T cells. **(E)** Number of DC (CD11c) or macrophage (FXIIIa) positive pixels overlapping with the body of CD4+ T cells (y axis) that harbor varying levels of HIV particles (x axis). Only images with at least 3 CD4+ T cells in each category (0, 1, 2-3, >=4 HIV particles) were selected for the analysis which consisted of 11 images across 5 donors. A Wilcoxon signed-rank test was performed to compare levels of DC or Mac pixel overlap between CD4+ T cells with varying levels of HIV. A Wilcoxon rank-sum test was performed to compare differences in the magnitude of membrane overlap between CD4+ T cells of varying HIV load. **(F)** Co-culture assay. Following cell sorting, human colorectum CD4+ T cells were plated alone or with DCs or macrophages at a 10:1 ratio (1 DC or macrophage per 10 CD4+ T cells). Cells were then exposed to HIVZ3678M at an MOI of 1 for 2h. Virus was washed from the culture media and cells were incubated for 72h and then assessed for dual p24 expression. A Wilcoxon rank-sum test was performed to compare the magnitude of the fold-change in p24 expression in co-cultures vs lone CD4+ T cell infections. **(G)** HIV transfer assay. Setup as in part F however PBMC-derived CD4+ T cells were added to the co-culture after first exposing DCs and macrophages to HIVZ3678M for 2h. Virus was washed from culture media, CD4+ T cells added and incubated for 72h followed by assessment of dual p24 expression. A Wilcoxon rank-sum test was performed to compare differences in the magnitude of the percentage of cells expressing p24.

To investigate potential HIV transfer events within HIV+ regions we devised an approach based on a modification of Neighbourhood Analysis. The number of HIV+ and HIV- cell neighbours were quantified for HIV+ cells. The locations of HIV+ cells, rather than all cells, were randomised to generate a null distribution of HIV+ cell neighbourhoods, against which the actual HIV+ cell neighbourhood was compared to determine significant interactions. Restricting randomisation to HIV+ cells acts as a control for the background effect of HIV on cell movement and allows for the measurement of a cell’s true interactions when it is HIV+. As HIV is present at the interface between cells during active viral transfer (Garcia et al., 2005; McDonald et al., 2003; Wang et al., 2007; Wang et al., 2008; Yu et al., 2008) HIV+:HIV- interactions were designated as ‘potential transfer events’ and HIV+:HIV+ interactions were designated as ‘likely transfer events’. Using this approach, we found that in the majority of images significant HIV+:HIV+ interactions occurred between pairs of target cells, including ‘DC:CD4+ T cell’ pairs in the LP and LAs, as well as ‘macrophage:CD4+ T cell’ pairs in the LP (**Figure 6C, top**). In each case the frequency of interactions for HIV+:HIV+ pairs was greater than the corresponding HIV+:HIV- pairs, indicating that the presence of HIV in both cells increases the likelihood of interaction. Interestingly, we also observed that cells of the same type preferred to interact with each other, whether or not they contained HIV, which has been noted in previous spatial studies analysing cell:cell interactions in tissue (Damond et al., 2019; Jackson et al., 2020; Keren et al., 2018).

The above approach is advantageous as it encompasses all events that influence the neighbourhood of HIV+ cells, such as migration to and sampling of HIV, as well as migration toward and the formation of clusters with other target cells. We also wished to quantify the propensity of a HIV+ cell to interact with and potentially transfer virus to a cell in its immediate neighbourhood. As such, we created a ‘local interaction score’, defined as the difference between the frequency of HIV+:HIV+ and HIV+:HIV- interactions (**Figure 6C bottom, STAR Methods**). Results showed that HIV+ DCs and macrophages had a substantial capacity to interact with local CD4+ T cells (∼50% increase in images showing significant interactions) (**Figure 6C, bottom, dotted boxes**). Interestingly, we also observed considerable local interactions between HIV+ DCs and macrophages. Importantly, HIV-binding induced no significant increase in local interactions with CD4- T cells in LP or ‘unknown’ cells in LAs. Representative images of DCs and macrophages interacting with CD4+ T cells, with HIV at their interface are shown in **Figure 6D** for HIV_BaL_ and **Figure S4C** for HIV_Z3678M_. We also observed DC-macrophage interactions with interfacing HIV using both HIV_BaL_ and HIV_Z3678M_ (**Figure S4D**).

To further characterise these interactions, we explored whether they had known characteristics of HIV transfer. In particular, *in vitro* studies using Scanning Electron Microscopy on blood-derived model DCs have shown that DCs use sheet-like membrane extensions to envelop CD4+ T cells which facilitates viral transfer and infection of CD4+ T cells (Felts et al., 2010). To investigate this, we measured membrane overlap of DCs and CD4+ T cells as a proxy for envelopment, hypothesising that CD4+ T cell HIV-load would correlate with increased membrane overlap. Indeed, increasing CD4+ T cell viral loads were associated with a significant increase in membrane overlap with DCs. Importantly this association was specific to DCs but not macrophages (**Figure 6E**). Additionally, DCs did not envelop CD4- T cells regardless of their (incidental) association with HIV (**Figure S4E**). This confirms the specificity of this phenomenon for ‘DC-CD4+ T cell’ interactions and controls for the possibility of the increased overlap being driven by DC migration toward HIV (**Figure 5A**), rather than HIV-containing DCs specifically seeking out CD4+ T cells.

Having observed that HIV+ DCs and macrophages preferentially interact with CD4+ T cells *in situ* we determined whether this led to increased viral replication in CD4+ T cells *ex vivo*. We sorted all colorectal HIV target cells and cocultured DCs or macrophages with autologous CD4+ T cells, prior to infection with a transmitted founder HIV strain and subsequent assessment of CD4+ T cell infection 72hrs later. We found that the presence of DCs or macrophages significantly enhanced infection of CD4+ T cells (**Figure 6F**). To determine whether enhancement was mediated by viral transfer we next infected DCs and macrophages prior to the addition of activated PBMC-derived CD4+ T cells. The results showed that both DCs and macrophages mediated HIV transfer to CD4+ T cells leading to increased viral replication but that this effect was significantly greater for DCs (**Figure 6G**).

All together, these results suggest that mucosal HIV entry induces its target cells to cluster together forming a community in which DCs and macrophages deliver virus to CD4+ T cells, facilitating infection of these cells within the mucosa itself.

## Discussion

In this study we developed a pipeline for multiplexed *in situ* quantification of initial host-pathogen transmission events and applied it to study human colorectal HIV transmission within 2h of exposure. This was made possible by a combination of RNAScope, CyCIF and image analysis algorithms to enable accurate quantification. In particular, RNAscope overcame issues of low signal-to-noise inherent in antibody and fluorophore-tagged HIV detection approaches (Campbell et al., 2007; Deleage et al., 2016; Eugenin and Berman, 2016; Prevedel et al., 2019) and, when combined with CyCIF, enabled quantitative comparison of HIV localisation to DCs, macrophages, CD4+ T cells and CD4- T cells (as a control) in four colorectal tissue compartments. Our AFid tool reduces false identification of HIV+ cells due to autofluorescence (Baharlou et al., 2020). Our segmentation approach allows cell overlap to outline full cell bodies. This enables accurate assignment of HIV particles to amorphous cells such as DCs and macrophages (**Figure S1**) whilst ensuring key cell:cell interactions aren’t missed as HIV can be transmitted between cells via membranous extensions (Eugenin et al., 2009; Nikolic et al., 2011). Finally, we employed spatial techniques to investigate cell:cell spread of HIV including our recently developed SpicyR algorithm (Canete et al., 2022), and a modified version of neighbourhood analysis (Schapiro et al., 2017). Together these approaches enabled us to dissect early transmission events *in situ* and provides a framework for *in situ* quantification of cellular microenvironment changes which could be applied to a range of disease and physiological settings.

The relative involvement of different HIV target cells in initial viral uptake is fundamental to understanding the determinants of transmission. However, this has not been previously described in intact primary human colorectal tissue. We reveal that HIV is first preferentially enriched in DCs and macrophages rather than CD4+ T cells, with DCs exhibiting the highest per-cell viral sampling capacity (**Figure 3**). Importantly, this was not replicable upon infection of *ex vivo* isolated rectal cells where macrophages were the dominant initial target cell, in agreement with previous work (Gurney et al., 2005). We postulate this difference is because the tissue microenvironment is a critical factor in HIV target cell migration and viral interactions, as illustrated in this study and others (Cavarelli et al., 2013; Ganor et al., 2010; Imle et al., 2019; Zhou et al., 2018). This reinforces the importance of *in situ* studies to accurately define initial host-pathogen interactions.

A key strength of this study is the analysis of tissue compartments (**Figures 4 and 5**) which revealed HIV enrichment in DCs and macrophages was specific to the mucosa and submucosa respectively, and that HIV preferentially localises to LAs. Mucosal DCs and submucosal macrophages steadily increased in density toward HIV particles, suggestive of migration to sites of HIV entry. LP DCs also appeared capable of crossing compartments to sample HIV, which has been observed for epithelium (Cavarelli et al., 2013), but which we report for the first time for LAs. As such, migration may be a key mechanism of early viral enrichment in these populations. This may be the function of one or more subsets of either DCs or macrophages, as subset-specific differences in both migratory capacity (Domanska et al., 2022) and HIV-binding (Bertram et al., 2019a; Rhodes et al., 2021) have been observed. HIV-binding itself appeared to influence cell location with LP HIV+ DCs and CD4+ T cells tending to locate near LAs. This could represent specific cell subsets in LA-adjacent regions, which are known to vary in murine studies of Peyer’s Patches (Bonnardel et al., 2015) but are unstudied in humans. Another possibility, at least for DCs, is that HIV-binding induces enhanced LA-directed migration and entry. This is supported by the increased DC density in HIV+ vs HIV- LAs and the preponderance of HIV+ DCs in these structures. Compartment-based analysis also revealed LAs are key HIV containing compartments within 2h of exposure to the virus. This may be due to delivery by HIV+ DCs or other cells in the adjacent LP, or via follicle-associated epithelium and resident M cells, which is a key site of entry for other enteric pathogens (Kobayashi et al., 2019). As LAs are enriched in rectal tissue (Langman and Rowland, 1986), contain abundant HIV target cells and are known sites of HIV persistence (Chun et al., 2008), rapid access to these structures may facilitate sustained infection and early reservoir formation within the mucosa. Indeed, naïve CD4+ T cells are enriched in LAs (Fenton et al., 2020) and are amenable to both viral transfer and activation by HIV+ DCs, which can promote latency (Cameron et al., 2010) and HIV-susceptible Th17 programs (Koh et al., 2020; Renault et al., 2022; Stieh et al., 2016). Entry into LAs but not LP was also associated with increased SM HIV levels, suggesting LAs as a possible conduit for HIV access to the underlying SM. This could be through direct passage as LAs traverse the muscularis mucosa barrier (Fenton et al., 2020) and HIV easily penetrates deep into LAs (**Figures S3H-J**). Alternatively, as HIV disseminates via lymphatics (Deleage et al., 2019), it may use the extensive LA lymphatic network (Fenton et al., 2020) to gain access to the SM, through which mucosal lymphatics drain (Unthank and Bohlen, 1988).

CD4+ T cells are the major initial targets of HIV integration and productive infection (Gupta et al., 2002; Maric et al., 2021; Stieh et al., 2016). However, the early events leading to infection remain poorly understood. DCs and macrophages facilitate HIV transfer and enhanced CD4+ T cell infection *in vitro*, however *in situ* characterisation using human tissue is lacking and mostly qualitative (Vine et al., 2022). Using our SpicyR algorithm and temporal inference we show that after HIV penetrates the EP, HIV target cells appear to move away from the EP and form clusters consisting of DCs, macrophages and CD4+ T cells (**Figure 6**). This may occur temporally as our data suggests macrophages migrate away from the EP first, followed by CD4+ T cells. Importantly, this was specific to HIV target cells and creates an ideal setting for viral transfer between cell types. Indeed, using a modified version of neighbourhood analysis, we show that upon HIV binding, DCs and macrophages preferentially cluster with CD4+ T cells, with virus present at the cellular interface. Interestingly, CD4+ T cell HIV load was tightly correlated with their physical overlap with DCs, suggesting that the early presence of high viral loads in CD4+ T cells may be dependent on their interactions with DCs. Indeed, infection of rectal tissue-derived co-cultures confirmed viral transfer and enhanced infection of CD4+ T cells by both DCs and macrophages, in agreement with other studies (Bertram et al., 2019a; Nasr et al., 2014; Rhodes et al., 2021). This provides the strongest evidence to date that viral transfer to CD4+ T cells occurs as early as 2h within the mucosa.

In summary this study uniquely contributes quantitative *in situ* data on the initial events of HIV transmission in intact human mucosal explant tissue. Although some of the findings of this study are circumstantial, we believe they give rise to important hypotheses regarding HIV transmission. Particularly, the role of niche-specific cell subsets, the drivers of cell recruitment and cell:cell interactions at sites of HIV entry, and the possibility of LAs as key sites of early viral amplification, persistence and extra-mucosal dissemination. Formal proof of early viral trafficking events would require 3D live cell imaging of intact human explants during viral invasion, which is challenging if not impossible with current technologies. However, we anticipate that recent advancements in high parameter imaging modalities (Baharlou et al., 2019; Lewis et al., 2021), animal models and human organoid systems (Kim et al., 2020) will allow unprecedented insight regarding the early determinants of HIV transmission. The approach and results presented here provide a foundation for such future studies which have the ability to inform prophylactic interventions or the design of a mucosal vaccine.

## Supplemental Information

This article includes 4 supplementary figures.

## Acknowledgements

This work was funded by National Health and Medical Research Council Ideas grant (GNT1181482). The authors would like to acknowledge the Westmead Cell Imaging and Flow Cytometry Core Facilities, supported by the Westmead Institute, Westmead Research Hub, Cancer Institute New South Wales.

## Author Contributions

Conceptualization: H.B., A.N.H

Methodology: H.B., A.N.H., N.C., E.E.V., K.M.B., J.W.R.

Experimental Investigation: H.B., E.E.V., D.Y.

Computational Analysis: H.B., N.C., K.H., E.P.

Novel Reagents and Samples: J.T., G.C., N.P., G.C., F.R., A.D.R., M.P.G

Visualization: H.B., K.H., E.E.V.

Writing: H.B., A.N.H.

Funding Acquisition: A.N.H., A.L.C.

Intellectual Input and Critical Revisions: H.B., A.N.H., A.L.C., E.E.V., K.J.S., K.M.B., N.N., S.N.B., J.D.E., M.A.H.

All authors read and approved of the manuscript.

## Declaration of Interests

The authors declare no competing interests.

## STAR Methods

### Key resources table

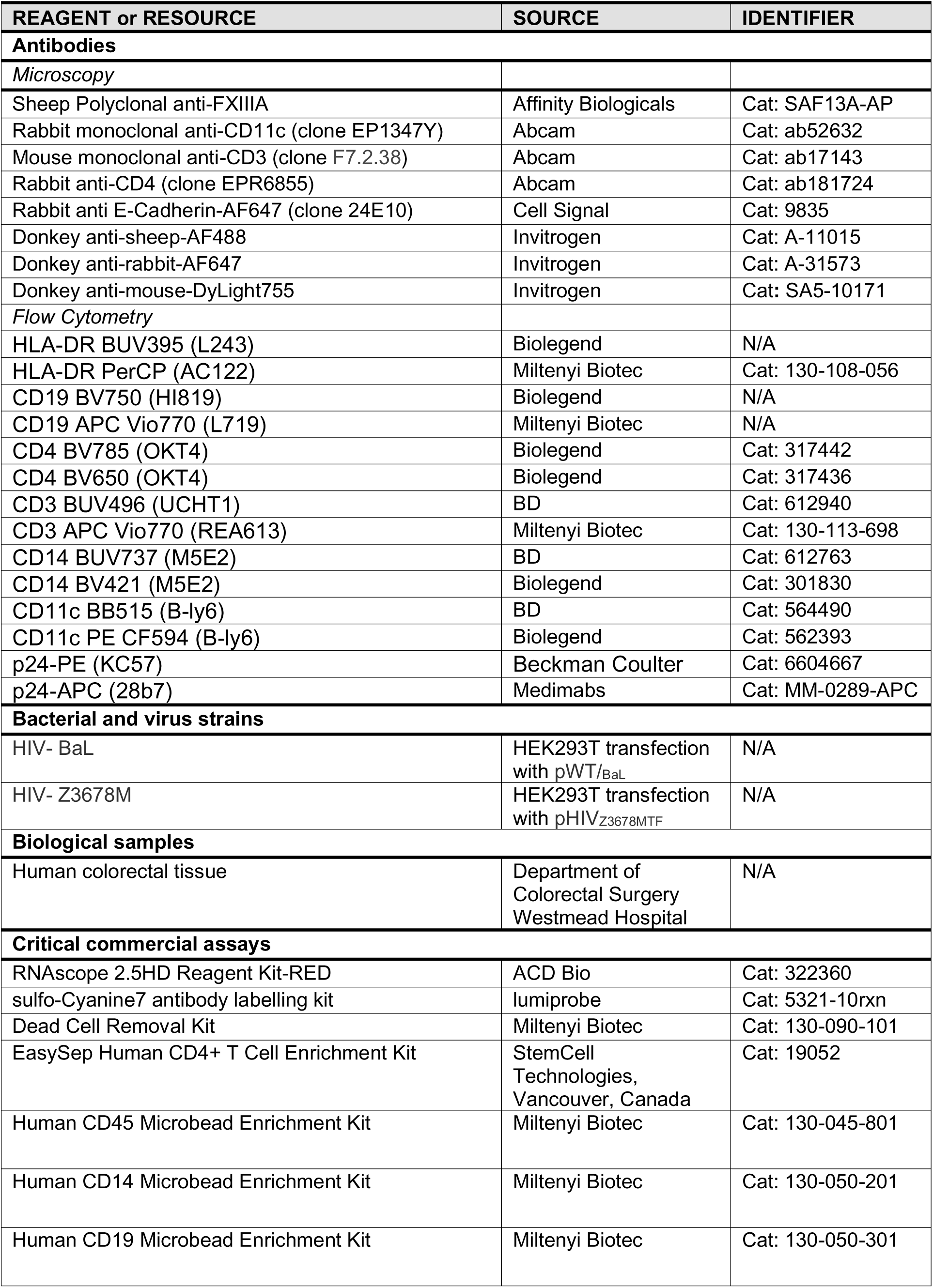
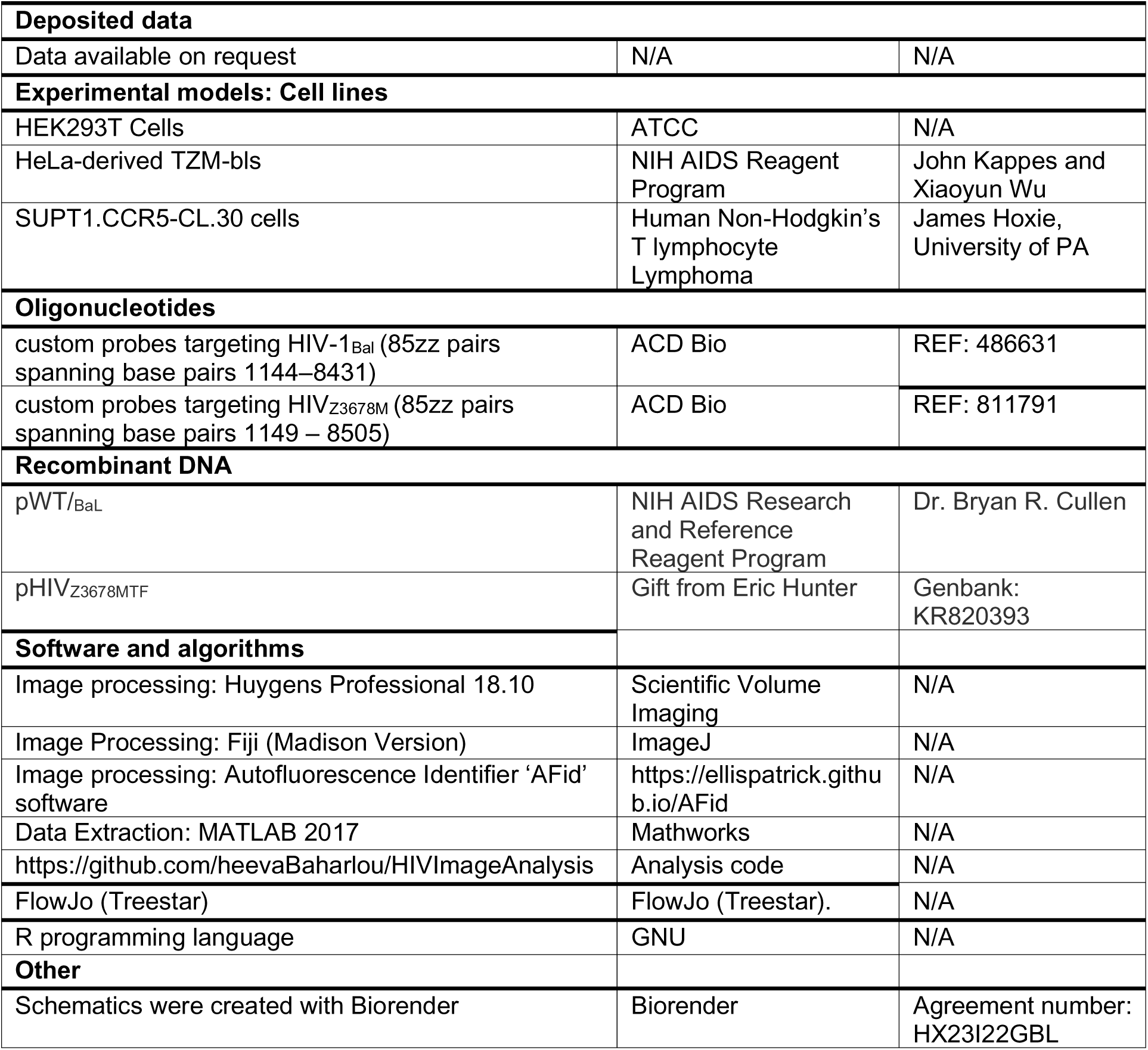

### Contact for Reagent and Resource Sharing

Further information and requests for resources and reagents should be directed to and will be fulfilled by the lead contact, Andrew Harman.

### Experimental Model and Subject Details

#### Human subjects

Healthy human colorectal tissue was obtained within 15 minutes of resection from patients undergoing surgical intervention for diverticulitis or colorectal cancer. Only healthy tissue distal to the site of disease process were used for this study. This study was approved by the Western Sydney Local Area Health District (WSLHD) Human Research Ethics Committee (HREC); reference number HREC/2013/8/4.4(3777) AU RED HREC/13/WMEAD/232. Written consent was obtained from all donors.

#### Cell lines

Both human embryonic kidney-derived 293T (HEK293T Cells) and HeLa-derived TZM-bls were cultured in Dulbecco’s Modified Eagle Medium (Lonza) with 10% Foetal Calf Serum (FCS) (Lonza) (DMEM10) at 37°C/5% CO_2_ and passaged using TrypLE express (Gibco) dissociation at a 1:10 dilution three times a week and 1:12 dilution twice a week, respectively. SUPT1.CCR5-CL.30 cells were maintained in RPMI (Lonza) with 10% FCS (RF10) at 37°C/5% CO_2_ and passaged at a 1:10 dilution twice per week.

### Method Details

#### HIV-1 virus production

Lab-adapted (HIV_BaL_) or transmitted founder (HIV_Z3678M_) strains were produced by transfection using a previously described protocol (Bertram et al., 2019a). 1.6×10^7^ HEK293T cells were seeded in a T150 flask (Becton Dickinson, Franklin Lakes,New Jersey, USA) and transfected with 80ug of pWT/_BaL_ or pHIV_Z3678MTF_ plasmid DNA. The following reaction mixture (all from Sigma-Aldrich) was prepared separately and added in addition to the plasmid DNA: 10μL 0.15M Na_2_HPO_4_ (pH 7.1), 128uL 2M CaCl_2_, 1 mL Hepes-buffered saline (280 mM NaCl, 50 mM HEPES, pH 7.1), 1 mL (1 mM Tris, 0.1 mM EDTA, pH 8.0), all diluted in 15 mL DMEM10 (Lonza). Culture media was replaced the next day with DMEM10 and cells cultured for a further 2 days, after which media was collected, centrifuged (1600g, 20min) and the resultant supernatant concentrated (3000g, 20min) to 1ml using 300kDa filters (Vivaspin 20, Sartorius, Göttingen, Germany). High titre stocks for HIV_BaL_ were achieved by infection of the SUPT1 T cell line. Stocks were depleted of microvesicles by adding 18ml viral supernatant to 2ml CD45 magnetic beads (Miltenyi Biotech) for 2h prior to filtering through an LS column (Miltenyi Biotech). The resultant CD45- HIV_BaL_ supernatant as well as HIV_Z3678M_ were under-layed with 1ml of 20% sucrose and ultracentrifuged (100,000g, 4°C, 1.5h, Beckman Optima XL-100 K, 70Ti rotor) to further concentrate viral stocks. This method yielded viral titres between 10^8^–10^9^, as measured by 50% tissue culture infective dose (TCID_50_)/mL on TZM-bl cells by LTR β-galactosidase reporter gene expression following one round of infection. Briefly, serial dilutions of viral stocks were performed on plated TZM-bls (37°C, 3 days), followed by media removal, addition of 50ul X-gal solution and incubation for 1h at 37°C. Wells were then diluted with 50ul of 4% PFA and incubated for a further 20min at room temperature (RT), followed by solution removal and EliSPOT imaging. The Spearman & Kärber algorithm was used for TCID_50_ measurements. Virus aliquots were stored at 80°C. Endotoxins (*Limulus* amebocyte lysate assay; Sigma), TNF-α, IFN-α, and IFN-β (Enzyme-linked immunosorbent assay (ELISA) were all below the limit of detection.

#### HIV explant infection

Human colorectal tissue was obtained within 15 minutes of surgical resection. Underlying fat and mesentery were removed using a scalpel and forceps and tissue spread out in a petri dish with the mucosal surface face-up. Gel-foam sponges (Pfizer) were cut into 1cm^2^ pieces (one for each explant), placed in the well of a 24-well plate and soaked in culture media consisting of 10μM HEPES (Gibco), non-essential amino acids (Gibco), 1 mM sodium pyruvate (Gibco), 50μM 2-Mercaptoethanol (Gibco), 10ug/ml Gentamycin (Gibco), 10% FCS (Lonza), all diluted in RPMI-1640 (Lonza). Here after this is referred to as ‘DC Culture Media’. Sponges were left to soak whilst tissue was further processed. As previously described (Doyle et al., 2021; Fenton et al., 2020) a dissection light microscope was used to select appropriate placement of cloning cylinders for the topical application of HIV. Appropriate areas were defined as containing visible lymphoid aggregates and being free of any signs of trauma. Once an appropriate area was located, an 8mm cloning cylinders (Sigma-Aldrich) was lightly coated on one side with histoacryl surgical glue (B Braun) using a fine paint brush and carefully placed over the region. 500ul of PBS was then added to the cloning cylinder to prevent the tissue surface from drying out and to hasten setting of the surgical glue. This step is critical as excess glue can slowly spread over time from the cylinder edge and cover the tissue surface thus preventing viral entry. Once all cylinders were placed, tissue within cylinders were checked again to ensure they were free of glue. 1 or 2 selected regions did not have cylinders placed but were resected with a scalpel as a 1cm^2^ area and fixed in 4% PFA (diluted in PBS) (electron microscopy sciences) for 18h. These samples were used to analyse target cell distribution in fresh uninfected colorectal tissue (**Figure 2**). The petri dish was then filled with PBS so as to just cover its surface, which reduced friction and the likelihood of displacement of the cylinders during cutting and lifting of explants. Soaked gel-foam sponges were then distributed across the 24-well plate. The perimeter surrounding each cloning cylinder was then cut, after which forceps were used to lift explants and place them on the sponges. Wells were then filled with DC Culture Media to the level of the tissue. A final quality check for cloning cylinder sealing was performed by ensuring that the previously applied PBS was still present and at the same level across explants. Solutions were removed from all cylinders and either 100ul PBS (mock), or a TCID_50_ of 70,000 (diluted in 100ul PBS) of lab adapted (HIV_BaL_) or transmitted founder (HIV_Z3678M_) strains applied to the inner chamber of cloning cylinders for 2h at 37°C. A TCID_50_ of 70,000 was selected as it is comparable to peak levels of infectious virus found in semen during the acute stage (Pilcher et al., 2007). Supernatant from inner chamber, as well as the DC Culture Media below were collected and stored at 80°C for assessment of HIV leakage from cylinders during the culture period (**Figure S3P**). Explants were then washed x3 with 500ul PBS to remove excess HIV from the mucosal surface prior to cylinder removal and tissue fixation in 4% PFA (diluted in PBS) for 18h.

#### Tissue embedding and sectioning

After fixation, explants were removed and trimmed with a scalpel along the impression left by the cloning cylinder during the culture period. This ensured that only areas exposed to virus remained. Explants were kept in 70% ethanol prior to embedding in paraffin. Tissues were then embedded flat so that the mucosal surface faced away from the sectioning surface of the paraffin block. This ensured that the thin mucosal surface would not accidently be trimmed away whilst sectioning. 4µm sections were taken through the entire tissue, from the submucosal side to the end of mucosa, and placed on Superfrost Ultra Plus Adhesion Slides (Thermo Scientific) Sections were stored in the dark at RT until staining.

#### RNAScope

Detection of HIV RNA was performed using RNAscope as previously described (Bertram et al., 2019a; Deleage et al., 2016; Rhodes et al., 2021) using the RNAscope 2.5HD Reagent Kit-RED’ with custom probes targeting HIV-1_Bal_ or HIV_Z3678M_. In the protocol that follows, reagents included in the ‘RNAscope 2.5HD Reagent Kit-RED’ are indicated. Unless otherwise written, wash steps were for 2min on a rotator set to low and incubations were carried out in a hybridisation oven (HybEZ Hybridization System (220VAC), ACD Bio). 4µm paraffin sections were baked at 60°C for 1h and dewaxed by sequentially submerging slides in xylene (2×2min) and 100% ethanol (2×2min). Slides were air-dried and antigen retrieval performed for 20min at 95°C using a pH9 buffer (RNAscope kit) and a decloaking chamber (Biocare). Sections were washed in TBS (Amresco, Cat: 0788), then Milli-Q H_2_O, followed by dipping slides 3-5 times in 100% ethanol, leaving them to air dry and then encircling sections with a hydrophobic pen (RNAscope kit). Sections were then incubated with protease pre-treatment 3 (diluted 1:5 in PBS and kept ice-cold) (RNAscope kit) for 30min at 40°C. Sections were washed x2 in Milli-Q H_2_O and incubated with probes targeting HIV-1_Bal_ or HIV_Z3678M_ for 2h at 40°C. Sections were washed x2 in RNAscope wash buffer (RNAscope kit). Signal from probes was then amplified using Amps 1-6 (RNAscope kit) which were added in sequence with x2 washes in RNAscope wash buffer between Amps. Amp incubation times were as follows Amp 1 = 30 min at 40 °C; Amp 2 = 15 min at 40 °C; Amp 3 = 30 min at 40 °C; Amp 4 = 15 min at 40 °C; Amp 5 = 30 min at RT; Amp 6 = 15 min at RT. HIV RNA Signal was developed using Fast Red substrate made by mixing Red-B and Red-A (RNAscope kit) at a 1:75 ratio for 5min at RT, followed by washing in Milli-Q H_2_O then TBS.

#### Cyclic Immunofluorescence staining and image acquisition

Unless otherwise indicated all washes were 2×2min in TBS on rotator set to low and incubations were in a humidified chamber protected from light. Following RNAscope, sections were blocked for 30min at RT with blocking buffer (10% donkey serum (Sigma), 1% BSA (Sigma), 0.1% Saponin (Sigma), all diluted in TBS) and washed. Sections were incubated with sheep anti-FXIIIa and rabbit anti-CD11c antibodies (diluted in block buffer) overnight at 4°C, washed and donkey anti-sheep-AF488 (Invitrogen, Cat: A-11015) and donkey anti-rabbit-AF647 (Invitrogen, Cat: A-31573) antibodies added for 30min at RT. Sections were washed and further blocked for 30min at RT with block buffer with 10% rabbit serum (DAKO) to block excess binding sites from the donkey anti-rabbit antibodies. 0.5% PFA (diluted in PBS) was added for 15min at RT to fix blocking rabbit IgGs in place. Rabbit anti-CD4-Cy7 (Abcam, Cat: ab181724, conjugation with sulfo-Cyanine7 antibody labelling kit (lumiprobe, Cat: 5321-10rxn)) was then added overnight at RT and sections washed 3×5min in TBS. Sections were stained with 1ug/ml DAPI (diluted in TBS) (Thermo Scientific, Cat: 62248) for 3min at RT, washed then rinsed in Milli-Q water. Sections were mounted with SlowFade Diamond Antifade Mountant (Invitrogen, Cat: S36963) and cover- slipped (Menzel-Glaser 22×60 mm Coverslip, Thermo Scientific). Images were acquired as per the section below on ‘Image Acquisition’. After imaging, slides were submerged in TBS until coverslips dissociated. Sections were then washed and treated with bleach solution (5% H2O2 (Sigma) and 20mM NaOH (Sigma) diluted diluted in PBS) for 1h with light (15 watt, 2700k light bulb, 5cm above sample). Sections were checked under the microscope to ensure signal removal from all channels, and subsequently washed and incubated with blocking buffer with 10% rabbit serum for 15min at RT, followed by washing in TBS. Rabbit E-Cadherin-AF647 (Cell Signal, Cat: 9835) and mouse anti-CD3 (Abcam, Cat: ab17143) were added overnight at RT. Sections were washed 3×5min in TBS and donkey anti-mouse-DyLight755 antibody (Invitrogen, Cat: SA5-10171) then added for 30min at RT. After washing 3×5min in TBS and rinsing in Milli-Q water slides were again mounted with SlowFade Diamond Antifade Mountant and imaged as described below.

#### Image acquisition

Images were acquired with a VS120 Slide Scanner equipped with an ORCA-FLASH 4.0 VS: Scientific CMOS camera (Olympus) and VS-ASW 2.9 software used for image acquisition and file conversion from vsi to tiff format. The entire tissue area was imaged using a x20 objective (UPLSAPO 20X/NA 0.75, WD 0.6/CG Thickness 0.17) and select areas for representative images were acquired using a x40 objective (UPLSAPO 40X/NA 0.95, WD 0.18/CG Thickness 0.11–0.23). For x40 images, Z-stacks were acquired 3.5µm above and below the plane of focus with 0.5µm step sizes. Channels used include: DAPI (Ex 387/11–25nm; Em: 440/40–25nm), FITC (Ex: 485/20–25nm; Em: 525/30–25nm), TRITC (Ex: 560/25–25 nm; Em: 607/36–25nm), Cy5 (Ex: 650/13–25 nm; Em: 700/75–75nm) and Cy7 (Ex: 710/75nm, Em: 810/90nm). All channels were checked, and antibodies titrated beforehand, to ensure against signal-spill over between channels.

#### Tissue digestion

To perform HIV-uptake (**Figure 3F**) and co-culture/transfer assays (**Figures 6F-G**), we extracted HIV target cells from human colorectal tissue. Underlying fat and mesentery were removed using a scalpel and forceps and remaining tissue cut into ∼5mm^2^ pieces. Surface epithelium and mucus was stripped by two incubations in RPMI with 10% FCS, 0.3% DTT (Sigma) and 2mM EDTA (Sigma) (15min, 37°C). Tissue was washed in DPBS and underwent two incubations in 20ml RPMI with 0.3% Collagenase IV (Worthington) and 0.5% DNase (Sigma) (30min, 37°C) to liberate cells which were then passed through a 100µm cell strainer and washed twice in DPBS. Cells were resuspended in 35ml RPMI, under-layed with 15ml Ficoll-Paque (GE Healthcare) and centrifuged (400g, 20min, no brakes). Buffy coats were collected and washed twice in DPBS. Red Cell Lysis buffer (All Sigma: 150 mM ammonium chloride (v/v), 10mM potassium bicarbonate (v/v), 0.1 mM EDTA (v/v) in ddH20) was used to remove remaining red blood cells as per the manufacturer’s instructions.

#### HIV uptake assay

Following our tissue digestion protocol (see above), liberated cells underwent positive selection for CD45+ cells (Miltenyi Biotec). 5×10^5^ cells in 150ul of DC Culture Media were then treated with HIV_Z3678M_ (MOI=5, 2h, 37°C). Cells were then washed three times in DPBS, 200ul of DPBS, stained with 0.05ul FVS700 for 30min at 4°C, washed in FACS wash (1% FCS (v/v), 2 mM EDTA, 0.1% sodium azide (w/v) in PBS) and 10ul Brilliant Stain Buffer (BD) added. Cells were then incubated with an antibody panel for 30min at 4°C with the final volume made to 50ul. The antibody panel included Biolegend: 1ul HLA-DR BUV395 (L243), 1ul CD19 BV750 (HI819), 2.5ul CD4 BV786 (OKT4); BD: 5ul CD3 BUV496 (UCHT1), 2.5ul CD14 BUV737 (M5E2), 1.5ul CD11c BB515 (B-ly6). Cells were washed in FACS wash, permeabilised with 100ul Cytofix/Cytoperm (BD) for 20min at RT and washed in Perm Wash (1% FCS (v/v), 1% BSA (w/v), 0.1% saponin (w/v), 0.1% sodium azide (w/v) in PBS) Cells were resuspended in 50ul Perm Wash and underwent intracellular staining with antibodies Beckman Coulter: p24-PE (KC57) and Medimabs: p24-APC (28b7) for 30min at RT. Cells were again washed in Perm Wash and HIV expression assessed by dual p24 expression using an LSRFortessa (BD).

#### Target cell selection and sorting

Following tissue digestion (see above), liberated cells were positively selected for CD45+ cells as per the manufacturer’s instructions (EasySep Human CD45+ Cell Enrichment Kit, StemCell Technologies) using a QuadroMACS separator with LS columns. 2.5×10^6^ cells were resuspended in 200ul of DPBS, stained with 0.05ul FVS700 for 30min at 4°C, washed in FACS wash and 10ul Brilliant Stain Buffer added. Our antibody sort panel was added for 30min at 4°C with the final volume made to 50ul. The sort panel included Miltenyi: 2.5ul CD3 APC Vio770 (REA613), 2.5ul CD19 APC Vio770 (L719), 1ul HLA-DR PerCP (AC122); Biolegend: 2ul CD4 BV650 (OKT4); BD: 1.5ul CD11c PE CF594 (B-ly6), 2.5ul CD14 BV421 (M5E2). Cells were then washed twice in FACS wash and once in pre-sort buffer (BD). 1ml of pre-sort buffer was used to resuspend cells which were then filtered using a 100µm cell strainer just prior to sorting on either the BDInflux (BD) or BDAriaIII (BD) cell sorters. CD4+ T cells were defined as live CD3+CD4+, Dendritic Cells as live HLA-DR+CD3-CD19-CD14- CD11c+ and Macrophages as live HLA-DR+CD3-CD19-CD14+. Sorted cells were placed in FACS tubes with 500ul of DC Culture Media and kept at 4°C until co-culture/transfer assay setup as described in the sections below.

#### HIV co-culture assay

Sorted DCs, macrophages and CD4+ T cells were plated as follows: CD4+ T cells alone, DCs with CD4+ T cells (1:10 ratio), macrophages with CD4+ T Cells (1:10 ratio). Cultures were topped to 150ul with DC Culture Media and treated with HIVZ3678M (MOI=1, 2h, 37°C). An additional mock-treated CD4+ T cell culture was maintained as a control. Cells were then washed three times in DC Culture Media, resuspended in 200ul of DC Culture Media with 0.02% Normocin (InvivoGen) and cultured for 3 days at 37°C. HIV infection of CD4+ T cells was determined by p24 expression using flow cytometry as described in the section on ‘HIV assessment by Flow Cytometry’.

#### HIV Transfer assay

Sorted DCs and macrophages as well as activated PBMC-derived CD4+ T cells (see section below) were plated in individual wells in 150ul DC Culture media, then exposed to HIVZ3678M (MOI=1, 2h, 37°C). An additional mock-treated activated CD4+ T cell culture was maintained as a control. Cells were then washed three times in DC Culture Media and resuspended in 200ul of DC Culture Media with 0.02% Normocin. Activated CD4+ T cells were then added to DC and Macrophage cultures at a 2:1 ratio and cultured for 3 days at 37°C. HIV infection of CD4+ T cells was determined by p24 expression using flow cytometry as described in the section on ‘HIV assessment by Flow Cytometry’.

#### HIV assessment by Flow Cytometry

Cells cultured with HIV were washed in DPBS, resuspended in 200ul, stained with FVS700 for 20min at 4°C and washed with FACS wash. Cells were stained with Miltenyi: CD3 APC-Vio770 (REA613) for 30min at 4°C, washed twice in FACS wash, permeabilised with 100ul Cytofix/Cytoperm for 20min at RT and washed in Perm Wash. Cells were resuspended in 50ul Perm Wash and underwent intracellular staining with antibodies Beckman Coulter: 1ul p24-PE (KC57) and Medimabs: 1ul p24-APC (28b7) for 30min at RT. Cells were again washed in Perm Wash and HIV expression assessed by dual p24 expression using an LSRFortessa.

#### PBMC-derived CD4+ T cell isolation

Transfer assays (**Figure 6G**) were performed using allogenic activated CD4+ T cells selected from peripheral blood mononuclear cells (PBMCs). PBMCs were derived from leukoreduction system chambers (LRSC) (Australian Red Cross Blood Service), on the same day as platelet donation. LRSCs were diluted 1:5, distributed across Falcon tubes with 35ml in each tube, then under-layed with 15ml Ficoll-Paque and centrifuged (400g, 20min, no brakes). Buffy coats were collected and washed x2 in DPBS. Red Cell Lysis buffer was used to remove remaining red blood cells as per the manufacturer’s instructions. CD4+ T cells were selected using a CD4 selection kit (StemCell Technologies) and activated by culturing 1×10^6^ cells/ml for 3 days at 37°C in RPMI supplemented with 10% FCS, 5 mg/mL PHA (Sigma) and 150 IU/mL IL-2 (Peprotech). Cells were transferred to cryovials containing FCS with 10% DMSO, placed in a CoolCell (Corning) and stored at -80°C.

### Quantification and Statistical Analysis

<u>Image analysis pipeline</u>

#### Image deconvolution and registration

Huygens Professional 18.10 (Scientific Volume Imaging, The Netherlands, http://svi.nl) CMLE algorithm, with SNR: 20 and 40 iterations was used for deconvolution of both single plane images acquired at x20, and also x40 Z-stacks. Images were aligned using the ImageJ plugin multiStackReg vs1.45 with the DAPI channel serving as a reference for alignment.

#### Autofluorescence removal

Colorectal tissue is prone to autofluorescence from many sources such as red blood cells, blood vessels, apoptotic cells, intrinsically autofluorescent cells etc. Our early analyses showed this substantially interfered with cell phenotyping and we could not remove autofluorescence using commercial quenching kits without significantly reducing our staining intensity. As such we developed ‘Autofluorescence Identifier’ (AFid) which analyses pixels from two input fluorescent channels and outlines autofluorescent objects (Baharlou et al., 2020). Pairs of channels compared were FXIIIA (on FITC) vs HIV RNA (on Texas Red) and HIV RNA vs CD11c. Autofluorescence masks were ‘OR’ combined and the resultant mask was used to exclude autofluorescent pixels (values set to 0) from images during data extraction as describted below. The input parameters for AFid were as follows:

Threshold: Niblack

Min Area = 20 pixels

Max Area = 100000 pixels

Sigma = 2 pixels

Correlation cut off = 0.6

Number of clusters = 1

Max Value to automate k = 0

Glow Removal = Yes

Expansion Sensitivity = 20 pixels

*Above values were for image resolutions of 3 pixels per µm.

#### HIV spot segmentation

HIV RNA particles were segmented using a custom MATLAB script based on a previously described spot counting algorithm (Battich et al., 2013). First, a manual threshold of the HIV RNA channel was set to approximate areas of HIV stain. The IdentifySpots2D function by Battich et al. was then used to identify the number of spots. The detection threshold was set to 0.01 and deblending steps was set to 2. Identified spots were excluded if they did not overlap with the manually generated threshold mask in the first step.

#### Cell Segmentation and Classification

Single cell segmentation was performed using a customised implementation of CellProfiler (Carpenter et al., 2006) in MATLAB (nucleiSegment.m, segRun.m function). Nuclei segmentation was performed by applying a local otsu filter to threshold the DAPI image, and applying object based watershed to identify boundaries. Objects with diameter between 3.3-16.7µm were kept. Masks of the CD3, CD11c and FXIIIa images were obtained by Gaussian blurring each image (sigma = 1.5) and performing a manual threshold to capture the full membrane. The nuclei is then classified based on the percentage overlap of each nuclei object with each membrane mask. A cell is classified as a T cell if the overlap with CD3 is >20%, classified as a DC if the overlap with CD11c is >20%, and classified as a macrophage if the overlap with FXIIIa is >40%. Finally, three separate labelled cell masks are obtained for each cell type by expanding the nuclei to fill the membrane one pixel at a time (expandNucleus.m). For example, a nuclei identified as a macrophage is expanded into the FXIIIa mask (**Sup Fig 1C**) space.

#### Tissue Compartment Segmentation

A manual threshold of the E-Cadherin stain was determined and nuclei (as segmented in the above section) belonging to this compartment were extracted using the BinaryReconstruct function in the ‘Morphology’ package in Fiji. The E-Cadherin and nuclei masks were then combined. The submucosa and lymphoid aggregates were manually outlined in Fiji and masks generated. A mask of the whole tissue was then generated with a manually determined threshold. Subtracting the epithelium, lymphoid aggregates and submucosa from the whole tissue mask provided a mask of the lamina propria. All masks were combined into a single image stack and assigned unique pixel values (eg, all epithelial pixels = 1, lamina propria pixels = 2 etc) so that they could be thresholded to extract data from each compartment.

#### Data Extraction in Matlab

Data extraction was performed in MATLAB (toRunNoNeighbrs.m, to RunNeighbrs.m). The mean marker expression and number of HIV particles was identified. From the compartment masks, a cell was considered to be part of that compartment if the overlap with the compartment mask was >25%. Distances from the compartments and HIV were obtained by creating a distance map from these objects and measuring the minimum value of the distance map within each cell. For cells within the LAs, the distance from the LA border was also measured in a similar approach. Neighbours were generated using the approach described in (Schapiro et al., 2017). Finally, using the cell masks, the overlap between cells were also identified. All data were exported into a single csv file with rows as individual cells and columns as cell features such CD4 expression, HIV particle number, distance from LAs etc.

#### Analysis in R

All image analysis for this study was performed in R using the csv spreadsheet of cells and their features, generated as described in the previous section. Procedures for statistical analysis generating the results in this paper are described in detail in the associated figure legends and links to the source code are provided in the Key Resources Table.

#### Location Interaction Score

We created a ‘local interaction score’ defined as the difference between the frequency of ‘HIV+:HIV+’ interactions (‘likely transfer events’) and ‘HIV+:HIV-‘ interactions (‘potential ‘transfer events’) measured by neighbourhood analysis. The ‘potential transfer events’ by definition incorporate all conditions leading up to a transfer event which includes the steady-state likelihood of two cells interacting and any HIV-induced effects on the movement of either cell just prior to a potential transfer. As such, if we take the difference in the frequency of significant interactions between likely and potential transfer events, we can remove the background interaction effect and directly measure the propensity of a HIV+ cell to transfer virus to another specific cell type in its immediate neighbourhood.

---

**Figure S1:**
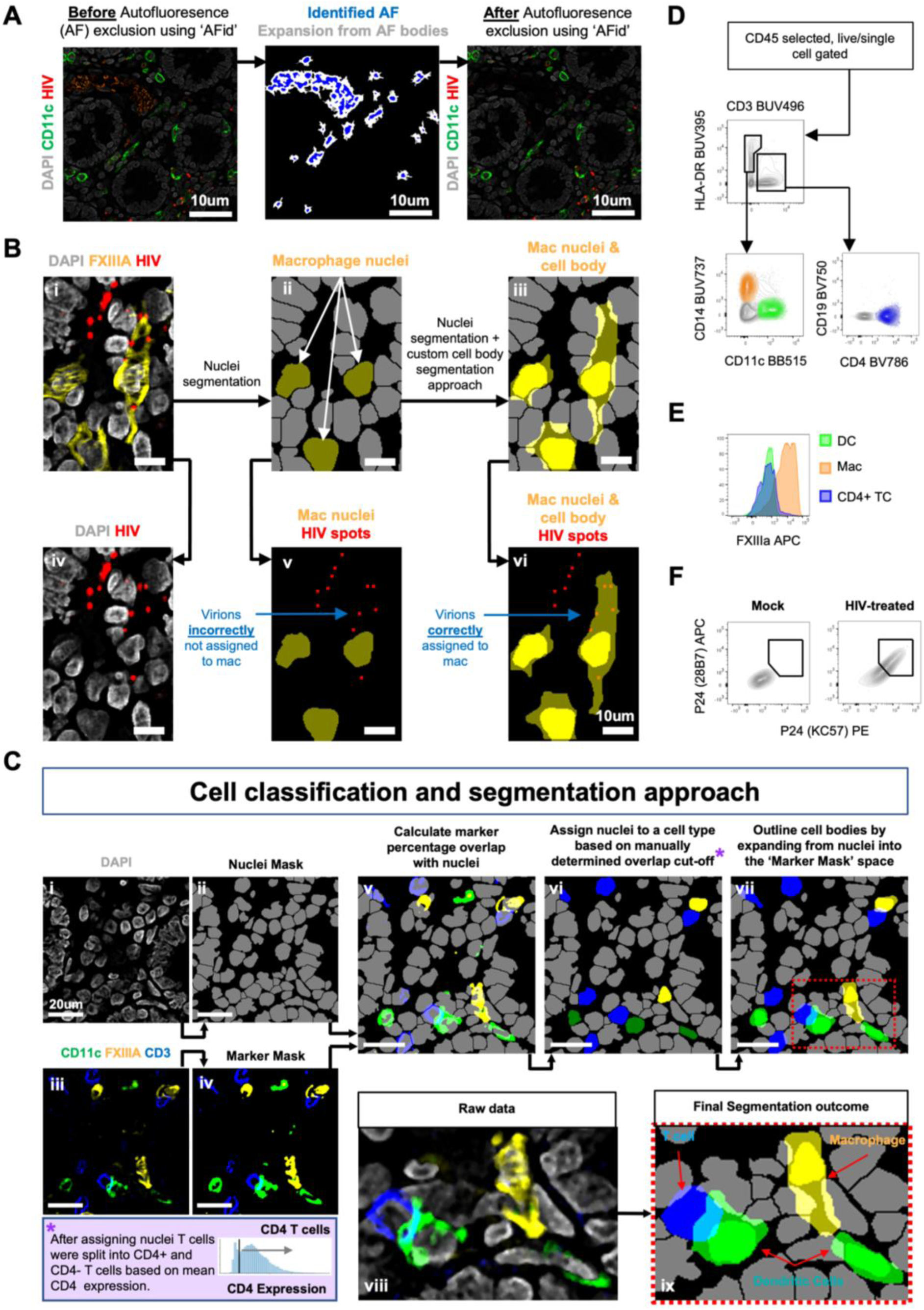
Cell Segmentation method (related to Figure 1) **(A)** Images before (left) and after (right) autofluorescence removal using AFid. The middle image shows the mask of the identified autofluorescence with core autofluorescent bodies in blue and the expansion mask shown in white. The expansion mask flows out from the autofluorescent bodies to capture all autofluorescence and is designed with a halting condition whereby the detection of stromal background fluorescence or antibody derived signal halts the expansion. Note that the algorithm also detects low signal autofluorescence and so some of the outlined autofluorescent bodies in the middle panel are not easily visible by eye. Importantly none of these signals overlap with the CD11c or HIV signals in the image. **(B)** These images illustrate the necessity of the custom segmentation approach outlined in part C for accurately assigning HIV virions to the correct cell type. A macrophage sampling multiple HIV particles is shown (i), as is the inadequacy of nuclei segmentation alone (ii) for accurately assigning HIV particles to the macrophage (v). This is rectified via the cell boundary estimation incorporated into our segmentation method (iii) which allows for the accurate assignment of HIV particles to the macrophage. **(C)** Cell classification and segmentation approach. First, a mask of the nuclei in each image was created by performing thresholding on DAPI followed by a morphological watershed (i-ii). Masks of CD11c, FXIIIa and CD3 were then created via manual thresholding (iii-iv). DCs, macrophages and T cells were classified by measuring the percentage overlap of marker masks from iv with the segmented nuclei from ii (v). The threshold for classifying a cell type was determined visually and was typically around 20% overlap for membrane markers CD11c and CD3 and 40% for the intracellular marker FXIIIa (vi). Once classified we then split the nuclei mask into three separate masks, each containing the nuclei belonging to one of the 3 cell types. In each mask, we then outlined the cell body, using distance maps emanating from nuclei centers, and restricted to the area of the masks of cell-type defining markers from part iv. This was followed by a morphological watershed to separate touching cells. The result was 3 masks which estimate the full cell body for each cell type (vii). Notably, a pixel from a single coordinate across the masks can belong to multiple cell types as the expansion in part vii was performed on separate masks. This is reflective of the actual situation in situ whereby cells interact in 3D and so will exhibit overlap with one another when taking 2D image slices. T cells were classified as CD4+ and CD4- by a manually determined cut off for CD4 expression (bottom left box). The raw data and the final segmentation outcome are shown side by side (viii-ix). **(D)** Gating strategy for HIV target cell identification by Flow Cytometry (Rhodes et al., 2021). **(E)** Representative histograms of FXIIIa expression on DCs, macrophages and CD4+ T cells (n = 3), showing FXIIIa expression is specific to macrophages as defined in part D. **(F)** Cell types defined in part D were either treated with HIV_Z3678M_ or untreated (mock) and stained with two different clones of antibodies (clones 28B7 and KC57) targeting p24. Dual p24+ positive cells were defined as HIV+ cells as per (Bertram et al., 2019a; Rhodes et al., 2021).

**Figure S2:**
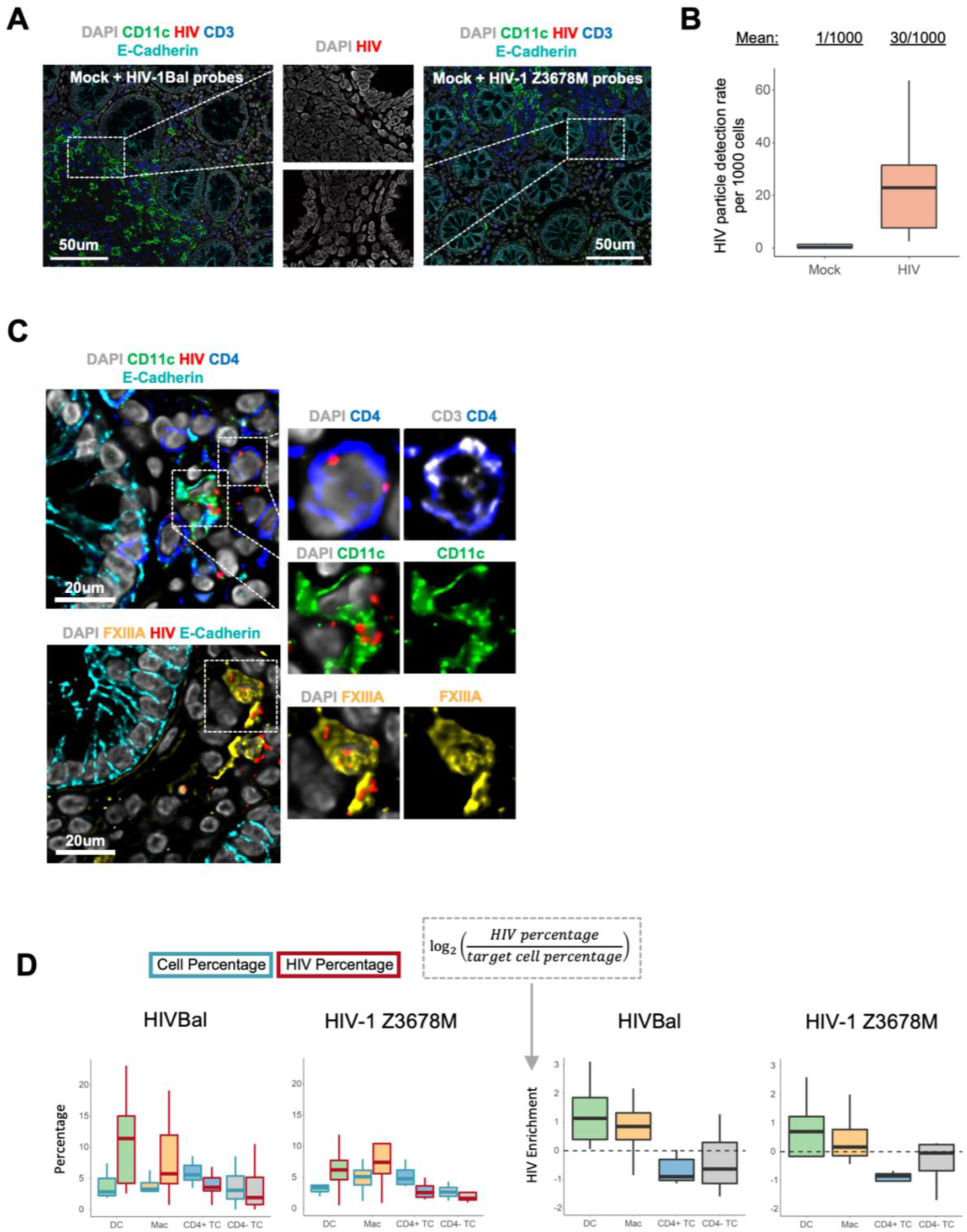
Assessment of interactions of HIV with colorectal target cells (related to Figure 3) **(A)** Images from mock (PBS treated) explants stained using probes against either HIV_Bal_ or HIV_Z3678M_. **(B)** HIV virions detected per 1000 cells in mock or HIV-treated samples stained using probes against either HIV_Bal_ or HIV_Z3678M_. **(C)** Representative images of colorectal target cells interacting with HIV_Z3678M_ particles. **(D)** Comparison of HIV uptake (HIV Percentage) relative to opportunity (Cell Percentage) across target cells and for two strains of HIV which were HIV_Bal_ and HIV_Z3678M_. Left: For each cell type their percentage among all cells (blue border) or the percentage of all HIV particles in those cells (red border) is shown. Right: HIV particle percentage normalized to the cell percentage, and log2 transformed. Data points represent individual donors, where the results from multiple images were averaged for each donor. HIV_Bal_: n = 11, HIV_Z3678M_: n = 4.

**Figure S3:**
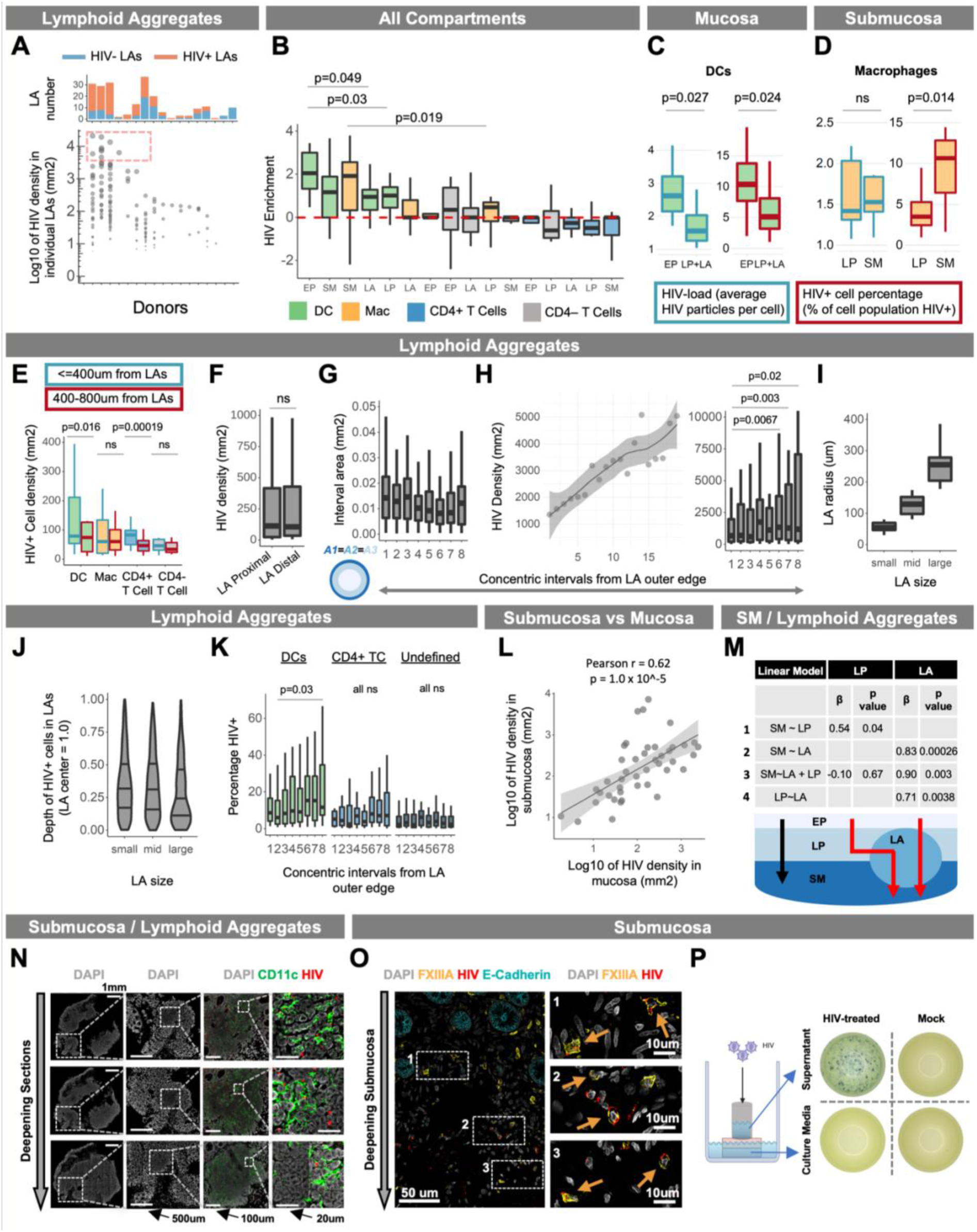
Differential HIV uptake across colorectal tissue compartments (related to Figure 4) **(A)** Density of HIV virions per mm^2^ in 215 LA images (y-axis) across 17 unique donors (x-axis). Each data point represents an individual LA from a given donor with dot size correlating to LA HIV density. The annotation above indicates the total number of LAs counted and is coloured by whether they are HIV+ (red) or HIV- (blue). Dotted box indicates a sample of LAs that were highly enriched with HIV (>4000 virions per mm^2^). **(B)** HIV enrichment in different cell types across tissue compartments. The HIV particle percentage was normalized to the cell percentage (data from Figure 4E), and log2 transformed. A Wilcoxon signed-rank test was performed to compare this HIV enrichment score across data points (specific cell types in specific tissue compartments). Data represent 15 explants from 12 donors, where the results from multiple images were averaged for each explant. **(C-D)** HIV-load (average virions per cell) or HIV+ cell percentage (percentage of cell population that is HIV+) was compared between ‘EP vs sub-EP’ compartments (LP + LA) for DCs and ‘LP vs SM’ for macrophages. Data represent individual donors where a donor was only included if at least 5 HIV+ cells were detected in both compartments for HIV-load measurements or 10 cells detected in both compartments for HIV+ cell percentage measurements. This was to allow for a fair comparison between compartments. A Wilcoxon rank-sum test was performed to compare measurements across compartments. **(E)** Density of HIV+ (>=1 particle) target cells and CD4- T cells in LA-proximal (<=400µm, blue border) vs -distal (400-800µm, red border) regions of LP. A distance of 400µm was chosen (instead of 200µm as in Figure 2D) in order to capture a larger pool of HIV+ cells, thus improving the reliability of comparisons. Data represents the ‘proximal’ and ‘distal’ LP from individual LAs. HIV+ cells for each cell type were compared between regions (Wilcoxon signed-rank test) only if each region contained at least 5 HIV+ cells, so as to allow for a fair comparison. **(F)** Density of HIV particles in LA-proximal (<=400µm) vs -distal (400-800µm) regions of LP. A Wilcoxon signed rank test was between LA-proximal and -distal regions of LP. **(G)** Area (mm^2^) of LA non-cumulative intervals. In order to achieve comparable areas for each interval we assumed LAs to be spherical and hence circular in 2D. We then calculated the radial edges of each interval from the outer edge using the formula: 1 – sqrt(1 – k/n) where k = {1,2…n-1} and n = max interval number. In this example n = 8. For a perfect circle this would derive intervals of equal area. **(H) Left:** Density of HIV+ cells in non-cumulative intervals from the outer edge of HIV+ LAs (x = 1) toward their center (x = 20). Results are shown as a LOESS curve of best fit to highlight cell density trends from outside LAs toward their center. **Right:** statistical comparisons (Wilcoxon signed rank test) of discrete intervals where LAs were split into 8 intervals instead of 20 in order to increase the number of cells measured per interval, thus reducing error and increasing the reliability of comparisons. **(I-J)** HIV+ cell depth in LAs (J) was measured in LAs categorized as small, medium or large (I). HIV+ cell depths were scaled such that ‘0 = LA outer edge’ and ‘1 = LA center’. LAs were assigned to each group by performing a quantile split, creating 3 even groups based on LA radius. Only LAs with at least 10 HIV+ cells were measured. Note that since area increases exponentially with increasing radius, ∼75% of the LA area exists half-way (y=0.5) toward the LA center. As such in this graph it appears as though HIV+ cell frequency is higher toward the LA edge, when in fact the opposite is true as shown in Figure S3G. **(K)** Percentage of LA DCs, CD4+ T cells or undefined cells (not a DC, Mac or T cell) that are HIV+ in non-cumulative intervals from the outer edge of HIV+ LAs (x = 1) toward their center (x = 8). A Wilcoxon signed rank test was performed between indicated intervals. **(L)** HIV virion density (per mm^2^ of DAPI) in SM vs mucosa (EP + LP + LA). Smoothed linear regression is shown with Pearson’s r and its associated p value. **(M)** Linear models of tissue-compartment HIV density to assess whether HIV entry into the SM is dependent on HIV entry into the LP or LAs. Linear models are shown in the leftmost column and the beta-weights and p-values for the independent variables (either LP or LA) are shown in the remaining columns. All variables correspond to the density of HIV virions in the compartment the variable is named after (i.e., SM = HIV virion density in SM). A schematic of the suggested pathways for viral entry into the SM is shown below the table, with the LA to SM entry suggested by the linear models shown by the red arrows. **(N)** Representative images showing HIV virions throughout various depths of a single LA from superficial sections in the mucosa (top row) to deeper sections where the LA has penetrated the submucosa (bottom row). **(O)** Representative images showing HIV+ Macs exist throughout various depths of the SM, including regions close to the LP (box 1) and deeper regions (box 3). **(P)** Representative images of TZM-bl assay results using either viral inoculum within cloning cylinders or the culture media in which the explants sit. Solutions were collected at the end of the viral culture period and then cultured with TZM-bls for 72h. Infected cells were visualized upon addition of X-gal substrate with each blue dot corresponding to an infected cell. All density measurements were performed per mm^2^ of DAPI for indicated regions.

**Figure S4:**
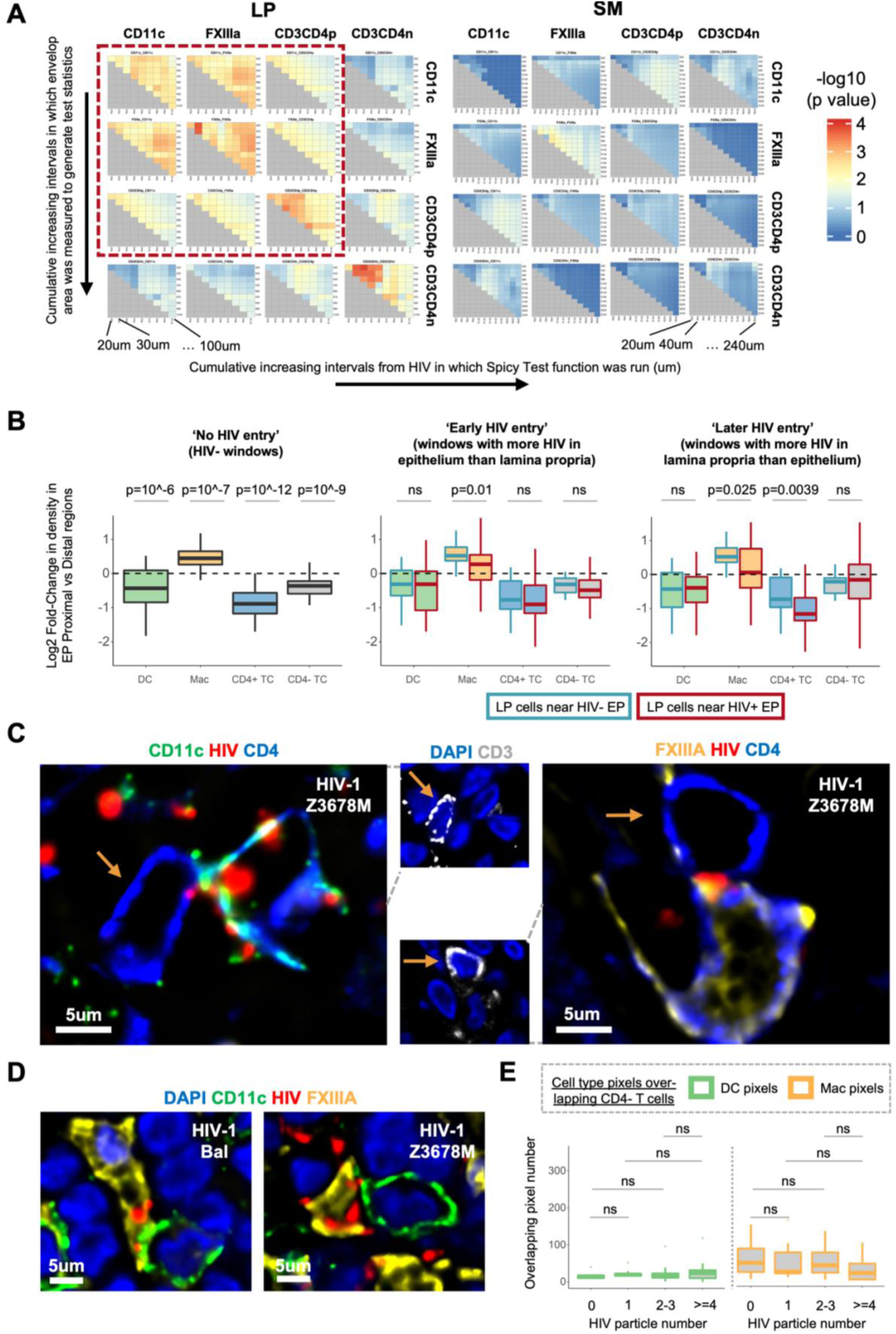
Dynamics of HIV-induced cell-cell interactions in situ (related to Figure 6) **(A)** SpicyR analysis as in Figure 6A with variation in radius (in which to measure cell-cell interactions) and HIV region distance cut-off (distance from HIV in which interactions were analysed). This analysis was run for LP and SM. The dotted red box encases HIV target cells in the lamina propria which can be seen to form significant clusters with most parameter settings. **(B)** LP target cell proximity to the EP in response to inferred stages of HIV entry. Images were divided into 100×100µm windows with each window classified as HIV- windows, ‘early’ (EP > LP HIV particle count) or ‘late’ (EP < LP HIV particle count) in terms of HIV entry. The log2 fold-change in LP target cell density in EP proximal (<=10µm from EP) vs distal regions (10-50µm from EP) was calculated for HIV- windows (left). For ‘early’ and ‘late’ stage windows the log2 fold-change was calculated from HIV- (blue border) and HIV+ EP (red border) separately. Only HIV- EP greater than 50µm from HIV+ EP were used. A Wilcoxon signed-rank test was used to compare ‘EP proximal vs EP distal fold-change in LP cells between HIV+ vs HIV- EP. Data from 40 images across 12 donors were used for this analysis. **(C)** Representative images of DCs and macrophages interacting with CD4+ T cells where HIV_Z3678M_ is present at the interface between the cells. For clarity, CD3 staining is shown in a separate image with a brown arrow pointing to CD3+CD4+ T cells. **(D)** Representative images of DCs and macrophages interacting with one another where HIV_Z3678M_ is present at the interface between these cells. **(E)** Number of DC (CD11c) or macrophage (FXIIIa) positive pixels overlapping with the body of CD4- T cells (y axis) that harbor varying levels of HIV particles (x axis). Only images with at least 3 CD4- T cells in each category (0, 1, 2-3, >=4 HIV particles) were selected for the analysis which consisted of 9 images across 5 donors. This comprised X images from Y donors. A Wilcoxon signed-rank test was performed to compare levels of DC or Mac pixel overlap between CD4- T cells with varying levels of HIV. A Wilcoxon rank-sum test was performed to compare differences in the magnitude of membrane overlap between CD4- T cells of varying HIV load. All density measurements were performed per mm^2^ of DAPI for indicated regions.

## Notes

### Competing Interest Statement

The authors have declared no competing interest.

