## Supplementary Figures for "An *in situ* quantitative map of initial human colorectal HIV transmission"

### Supplementary Figure 1

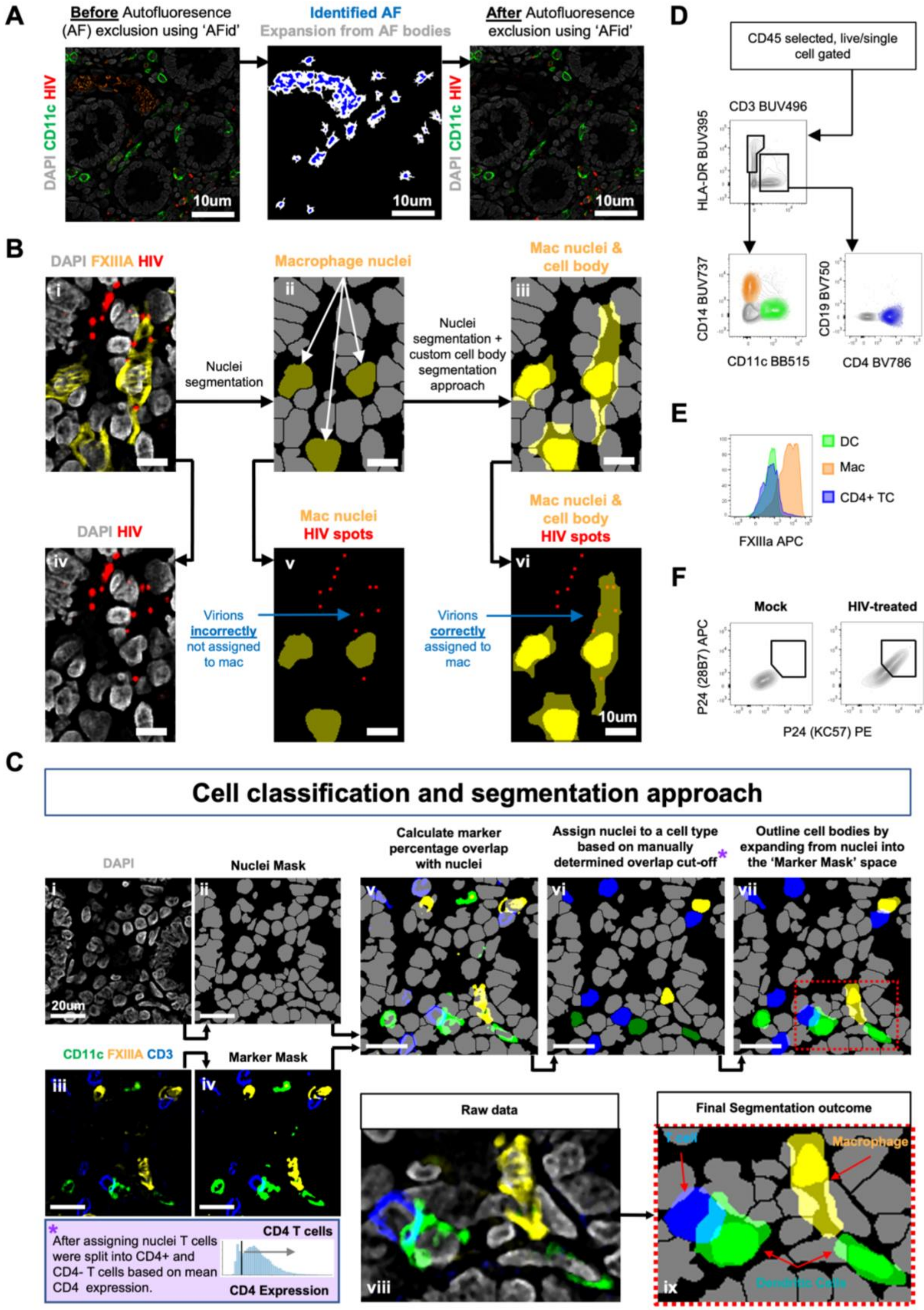

**Figure S1: Cell Segmentation method (related to Figure 1)**

**(A)** Images before (left) and after (right) autofluorescence removal using AFid. The middle image shows the mask of the identified autofluorescence with core autofluorescent bodies in blue and the expansion mask shown in white. The expansion mask flows out from the autofluorescent bodies to capture all autofluorescence and is designed with a halting condition whereby the detection of stromal background fluorescence or antibody derived signal halts the expansion. Note that the algorithm also detects low signal autofluorescence and so some of the outlined autofluorescent bodies in the middle panel are not easily visible by eye. Importantly none of these signals overlap with the CD11c or HIV signals in the image.

**(B)** These images illustrate the necessity of the custom segmentation approach outlined in part C for accurately assigning HIV virions to the correct cell type. A macrophage sampling multiple HIV particles is shown (i), as is the inadequacy of nuclei segmentation alone (ii) for accurately assigning HIV particles to the macrophage (v). This is rectified via the cell boundary estimation incorporated into our segmentation method (iii) which allows for the accurate assignment of HIV particles to the macrophage.

**(C)** Cell classification and segmentation approach. First, a mask of the nuclei in each image was created by performing thresholding on DAPI followed by a morphological watershed (i-ii). Masks of CD11c, FXIIIa and CD3 were then created via manual thresholding (iii-iv). DCs, macrophages and T cells were classified by measuring the percentage overlap of marker masks from iv with the segmented nuclei from ii (v). The threshold for classifying a cell type was determined visually and was typically around 20% overlap for membrane markers CD11c and CD3 and 40% for the intracellular marker FXIIIa (vi). Once classified we then split the nuclei mask into three separate masks, each containing the nuclei belonging to one of the 3 cell types. In each mask, we then outlined the cell body, using distance maps emanating from nuclei centers, and restricted to the area of the masks of cell-type defining markers from part iv. This was followed by a morphological watershed to separate touching cells. The result was 3 masks which estimate the full cell body for each cell type (vii). Notably, a pixel from a single coordinate across the masks can belong to multiple cell types as the expansion in part vii was performed on separate masks. This is reflective of the actual situation in situ whereby cells interact in 3D and so will exhibit overlap with one another when taking 2D image slices. T cells were classified as CD4+ and CD4- by a manually determined cut off for CD4 expression (bottom left box). The raw data and the final segmentation outcome are shown side by side (viii-ix).

**(D)** Gating strategy for HIV target cell identification by Flow Cytometry (Rhodes et al., 2021).

**(E)** Representative histograms of FXIIIa expression on DCs, macrophages and CD4+ T cells (n = 3), showing FXIIIa expression is specific to macrophages as defined in part D.

**(F)** Cell types defined in part D were either treated with $HIV_{Z3678M}$ or untreated (mock) and stained with two different clones of antibodies (clones 28B7 and KC57) targeting p24. Dual p24+ positive cells were defined as HIV+ cells as per (Bertram et al., 2019a; Rhodes et al., 2021).

### Figure S2

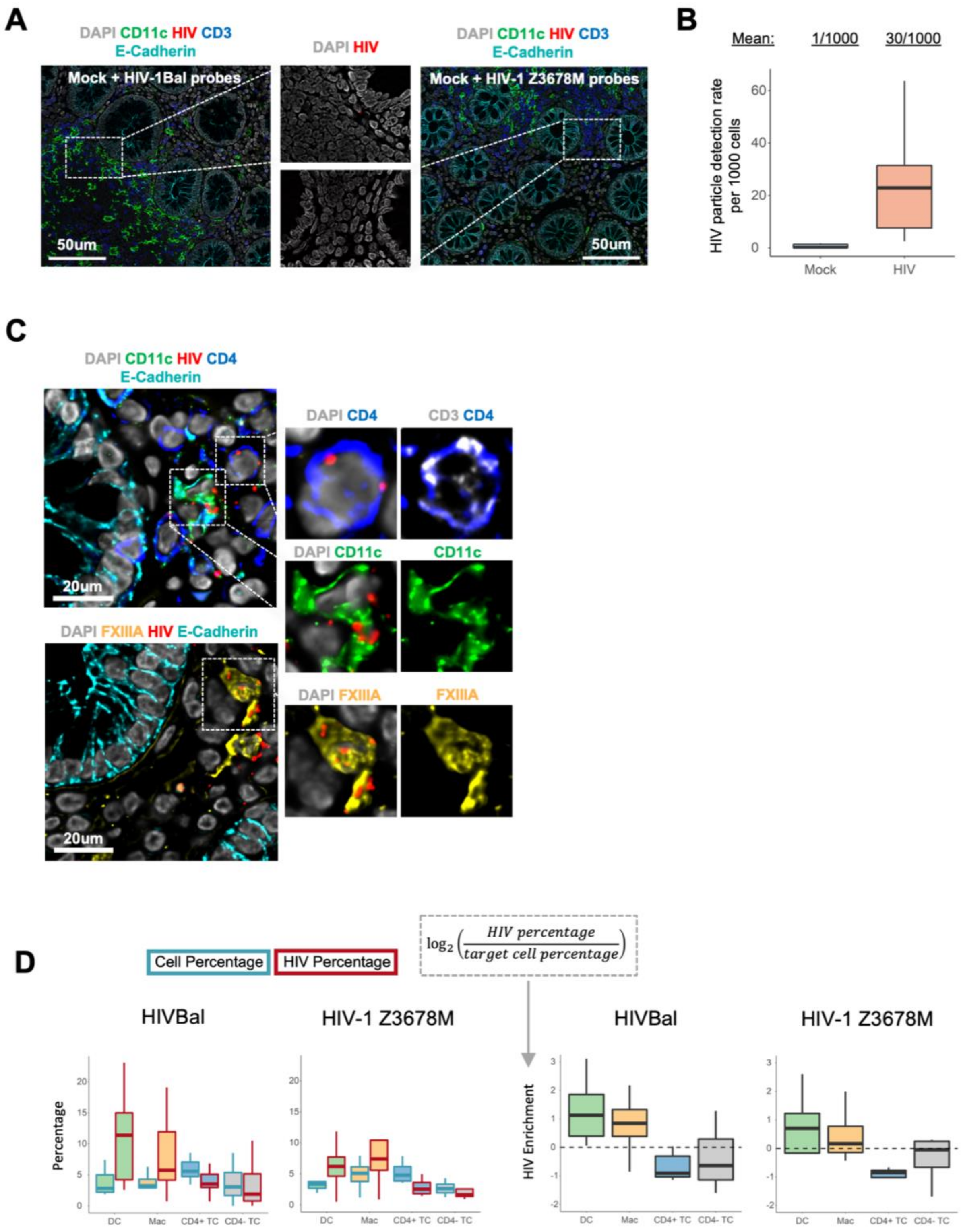
A
DAPI CD11c HIV CD3
E-Cadherin
Mock + HIV-1Bal probes
50um
DAPI HIV
Mock + HIV-1 Z3678M probes
B
Mean: 1/1000 30/1000
HIV particle detection rate per 1000 cells
0
20
40
60
Mock
HIV
C
DAPI CD11c HIV CD4
E-Cadherin
20um
DAPI CD4
CD3 CD4
DAPI CD11c
CD11c
DAPI FXIIIA HIV E-Cadherin
DAPI FXIIIA
FXIIIA
D
Cell Percentage
HIV Percentage
$\log_2\left(\frac{HIV\ percentage}{target\ cell\ percentage}\right)$
HIVBal
HIV-1 Z3678M
Percentage
HIV Enrichment
DC
Mac
CD4+ TC
CD4- TC

**Figure S2: Assessment of interactions of HIV with colorectal target cells (related to Figure 3)**

**(A)** Images from mock (PBS treated) explants stained using probes against either $HIV_{Bal}$ or $HIV_{Z3678M}$.

**(B)** HIV virions detected per 1000 cells in mock or HIV-treated samples stained using probes against either $HIV_{Bal}$ or $HIV_{Z3678M}$.

**(C)** Representative images of colorectal target cells interacting with $HIV_{Z3678M}$ particles.

**(D)** Comparison of HIV uptake (HIV Percentage) relative to opportunity (Cell Percentage) across target cells and for two strains of HIV which were $HIV_{Bal}$ and $HIV_{Z3678M}$. Left: For each cell type their percentage among all cells (blue border) or the percentage of all HIV particles in those cells (red border) is shown. Right: HIV particle percentage normalized to the cell percentage, and log2 transformed. Data points represent individual donors, where the results from multiple images were averaged for each donor. $HIV_{Bal}$: n = 11, $HIV_{Z3678M}$: n = 4.

### Figure S3

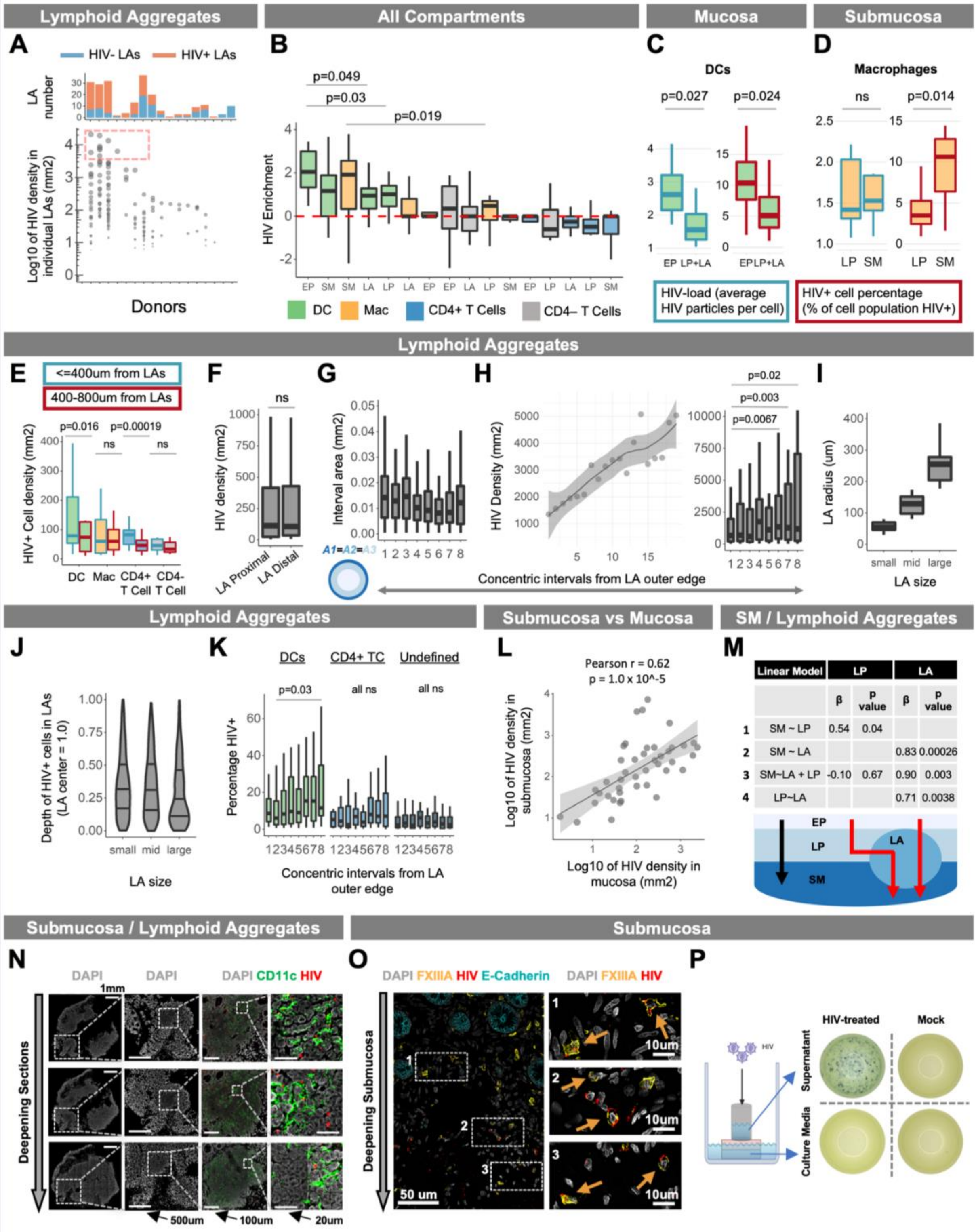


| Linear Model | LP | | LA | |
|---|---|---|---|---|
| | β | p value | β | p value |
| 1 SM ~ LP | 0.54 | 0.04 | | |
| 2 SM ~ LA | | | 0.83 | 0.00026 |
| 3 SM~LA + LP | -0.10 | 0.67 | 0.90 | 0.003 |
| 4 LP~LA | | | 0.71 | 0.0038 |

**Figure S3: Differential HIV uptake across colorectal tissue compartments (related to Figure 4)**

**(A)** Density of HIV virions per $mm^2$ in 215 LA images (y-axis) across 17 unique donors (x-axis). Each data point represents an individual LA from a given donor with dot size correlating to LA HIV density. The annotation above indicates the total number of LAs counted and is coloured by whether they are HIV+ (red) or HIV- (blue). Dotted box indicates a sample of LAs that were highly enriched with HIV (>4000 virions per $mm^2$).

**(B)** HIV enrichment in different cell types across tissue compartments. The HIV particle percentage was normalized to the cell percentage (data from Figure 4E), and log2 transformed. A Wilcoxon signed-rank test was performed to compare this HIV enrichment score across data points (specific cell types in specific tissue compartments). Data represent 15 explants from 12 donors, where the results from multiple images were averaged for each explant.

**(C-D)** HIV-load (average virions per cell) or HIV+ cell percentage (percentage of cell population that is HIV+) was compared between 'EP vs sub-EP' compartments (LP + LA) for DCs and 'LP vs SM' for macrophages. Data represent individual donors where a donor was only included if at least 5 HIV+ cells were detected in both compartments for HIV-load measurements or 10 cells detected in both compartments for HIV+ cell percentage measurements. This was to allow for a fair comparison between compartments. A Wilcoxon rank-sum test was performed to compare measurements across compartments.

**(E)** Density of HIV+ (>=1 particle) target cells and CD4- T cells in LA-proximal (<=400µm, blue border) vs -distal (400-800µm, red border) regions of LP. A distance of 400µm was chosen (instead of 200µm as in Figure 2D) in order to capture a larger pool of HIV+ cells, thus improving the reliability of comparisons. Data represents the 'proximal' and 'distal' LP from individual LAs. HIV+ cells for each cell type were compared between regions (Wilcoxon signed-rank test) only if each region contained at least 5 HIV+ cells, so as to allow for a fair comparison.

**(F)** Density of HIV particles in LA-proximal (<=400µm) vs -distal (400-800µm) regions of LP. A Wilcoxon signed rank test was between LA-proximal and -distal regions of LP.

**(G)** Area ($mm^2$) of LA non-cumulative intervals. In order to achieve comparable areas for each interval we assumed LAs to be spherical and hence circular in 2D. We then calculated the radial edges of each interval from the outer edge using the formula: $1 - \text{sqrt}(1 - k/n)$ where $k = \{1,2\ldots n-1\}$ and $n$ = max interval number. In this example $n = 8$. For a perfect circle this would derive intervals of equal area.

**(H) Left:** Density of HIV+ cells in non-cumulative intervals from the outer edge of HIV+ LAs (x = 1) toward their center (x = 20). Results are shown as a LOESS curve of best fit to highlight cell density trends from outside LAs toward their center. **Right:** statistical comparisons (Wilcoxon signed rank test) of discrete intervals where LAs were split into 8 intervals instead of 20 in order to increase the number of cells measured per interval, thus reducing error and increasing the reliability of comparisons.

**(I-J)** HIV+ cell depth in LAs (J) was measured in LAs categorized as small, medium or large (I). HIV+ cell depths were scaled such that '0 = LA outer edge' and '1 = LA center'. LAs were assigned to each group by performing a quantile split, creating 3 even groups based on LA radius. Only LAs with at least 10 HIV+ cells were measured. Note that since area increases exponentially with increasing radius, ~75% of the LA area exists half-way (y=0.5) toward the LA center. As such in this graph it appears as though HIV+ cell frequency is higher toward the LA edge, when in fact the opposite is true as shown in Figure S3G.

**(K)** Percentage of LA DCs, CD4+ T cells or undefined cells (not a DC, Mac or T cell) that are HIV+ in non-cumulative intervals from the outer edge of HIV+ LAs (x = 1) toward their center (x = 8). A Wilcoxon signed rank test was performed between indicated intervals.

**(L)** HIV virion density (per $mm^2$ of DAPI) in SM vs mucosa (EP + LP + LA). Smoothed linear regression is shown with Pearson's r and its associated p value.

**(M)** Linear models of tissue-compartment HIV density to assess whether HIV entry into the SM is dependent on HIV entry into the LP or LAs. Linear models are shown in the leftmost column and the beta-weights and p-values for the independent variables (either LP or LA) are shown in the remaining columns. All variables correspond to the density of HIV virions in the compartment the variable is named after (i.e., SM = HIV virion density in SM). A schematic of the suggested pathways for viral entry into the SM is shown below the table, with the LA to SM entry suggested by the linear models shown by the red arrows.

**(N)** Representative images showing HIV virions throughout various depths of a single LA from superficial sections in the mucosa (top row) to deeper sections where the LA has penetrated the submucosa (bottom row).

**(O)** Representative images showing HIV+ Macs exist throughout various depths of the SM, including regions close to the LP (box 1) and deeper regions (box 3).

**(P)** Representative images of TZM-bl assay results using either viral inoculum within cloning cylinders or the culture media in which the explants sit. Solutions were collected at the end of the viral culture period and then cultured with TZM-bls for 72h. Infected cells were visualized upon addition of X-gal substrate with each blue dot corresponding to an infected cell.

All density measurements were performed per $mm^2$ of DAPI for indicated regions.

### Supplementary Figure 4

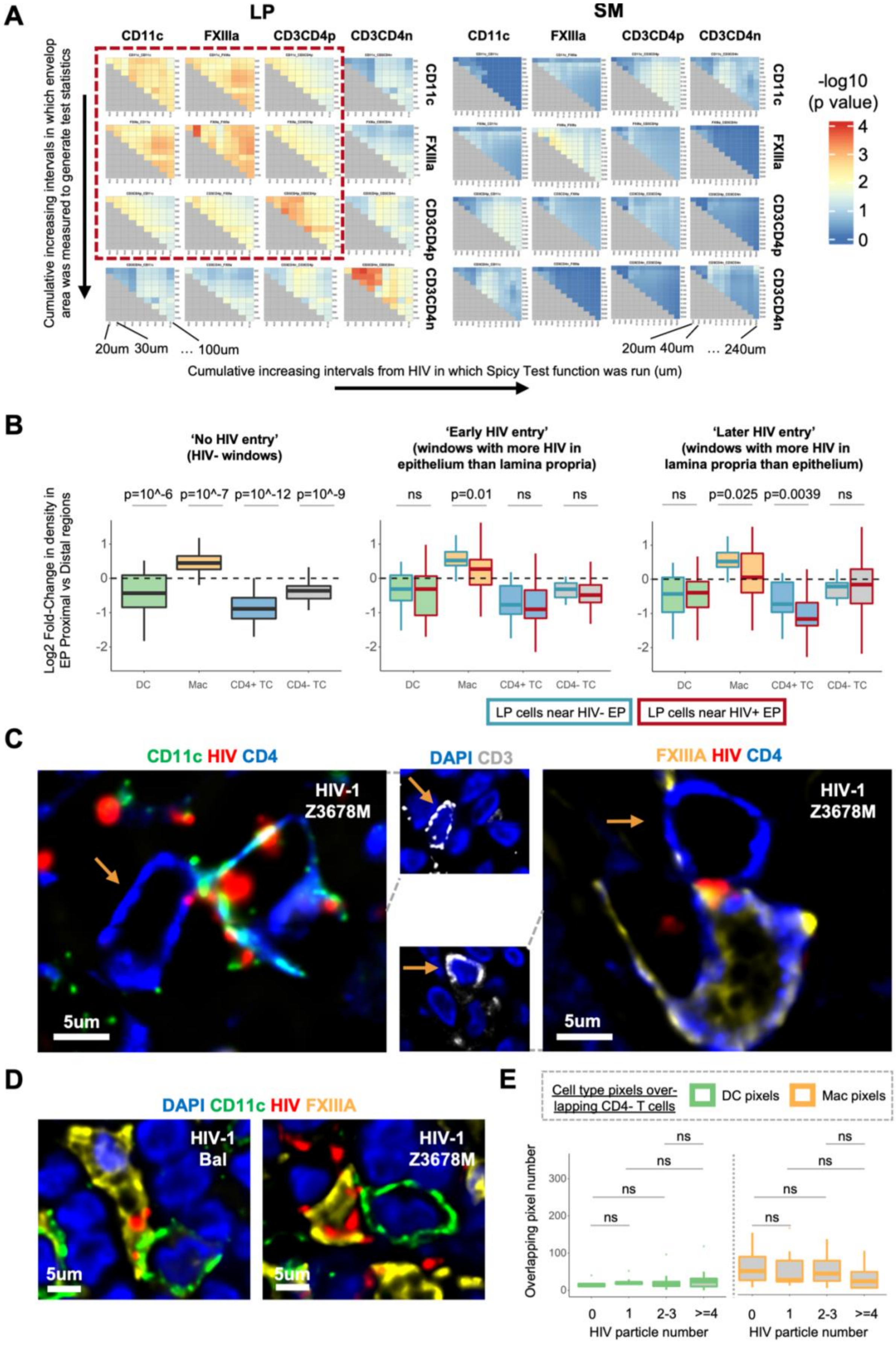

**Figure S4: Dynamics of HIV-induced cell-cell interactions in situ (related to Figure 6)**

**(A)** SpicyR analysis as in Figure 6A with variation in radius (in which to measure cell-cell interactions) and HIV region distance cut-off (distance from HIV in which interactions were analysed). This analysis was run for LP and SM. The dotted red box encases HIV target cells in the lamina propria which can be seen to form significant clusters with most parameter settings.

**(B)** LP target cell proximity to the EP in response to inferred stages of HIV entry. Images were divided into 100x100µm windows with each window classified as HIV- windows, 'early' (EP > LP HIV particle count) or 'late' (EP < LP HIV particle count) in terms of HIV entry. The log2 fold-change in LP target cell density in EP proximal (<=10µm from EP) vs distal regions (10-50µm from EP) was calculated for HIV- windows (left). For 'early' and 'late' stage windows the log2 fold-change was calculated from HIV- (blue border) and HIV+ EP (red border) separately. Only HIV- EP greater than 50µm from HIV+ EP were used. A Wilcoxon signed-rank test was used to compare 'EP proximal vs EP distal fold-change in LP cells between HIV+ vs HIV- EP. Data from 40 images across 12 donors were used for this analysis.

**(C)** Representative images of DCs and macrophages interacting with CD4+ T cells where $HIV_{Z3678M}$ is present at the interface between the cells. For clarity, CD3 staining is shown in a separate image with a brown arrow pointing to CD3+CD4+ T cells.

**(D)** Representative images of DCs and macrophages interacting with one another where $HIV_{Z3678M}$ is present at the interface between these cells.

**(E)** Number of DC (CD11c) or macrophage (FXIIIa) positive pixels overlapping with the body of CD4- T cells (y axis) that harbor varying levels of HIV particles (x axis). Only images with at least 3 CD4- T cells in each category (0, 1, 2-3, >=4 HIV particles) were selected for the analysis which consisted of 9 images across 5 donors. This comprised X images from Y donors. A Wilcoxon signed-rank test was performed to compare levels of DC or Mac pixel overlap between CD4- T cells with varying levels of HIV. A Wilcoxon rank-sum test was performed to compare differences in the magnitude of membrane overlap between CD4- T cells of varying HIV load.

All density measurements were performed per $mm^2$ of DAPI for indicated regions.
